# Postnatal Maturation of Dendritic Epidermal T Cells and Langerhans Cells Follows Distinct Differentiation Trajectories Independent of Microbiota

**DOI:** 10.64898/2026.04.05.716534

**Authors:** David Obwegs, Alexander Oschwald, Lara M. Koetter, Cylia Crisand, Sidney Doerr, Kerstin Bruder, Solveig Runge, Nisreen Ghanem, Vidmante Fuchs, Marleen Eckert, Julia Kolter, Daniel Erny, Marco Prinz, Susana Minguet, Wolfgang W. Schamel, Philipp Henneke, Stephan P. Rosshart, Katrin Kierdorf, Sagar

## Abstract

The mouse epidermis harbors two key resident immune populations—dendritic epidermal T cells (DETCs), a subset of invariant γδ T cells, and Langerhans cells (LCs), specialized tissue-resident macrophages—both of which play critical roles in immune surveillance, barrier integrity, and tissue homeostasis. While the fetal origin of both cell types has been defined, the cellular and molecular mechanisms that govern their postnatal fates following colonization of the epidermis around birth remain incompletely understood. Here, we present a combination of immunophenotyping- and transcriptome-resolved single-cell map of DETC and LC development in the mouse epidermis from late embryogenesis through adulthood. We delineate differentiation trajectories for both cell types, marked by distinct changes in morphology, proliferation, and transcriptional programming. Using mice deficient in γδ T cells, which lack canonical DETCs, we demonstrate that LCs develop independently of canonical DETCs likely due to the presence of αβDETCs. Moreover, analysis of germ-free mice and wildlings reveals that the postnatal development of both DETCs and LCs is independent of microbial colonization. Together, our findings define the core principles underlying the establishment of the mouse epidermal immune niche.

**GRAPHICAL ABSTRACT:** 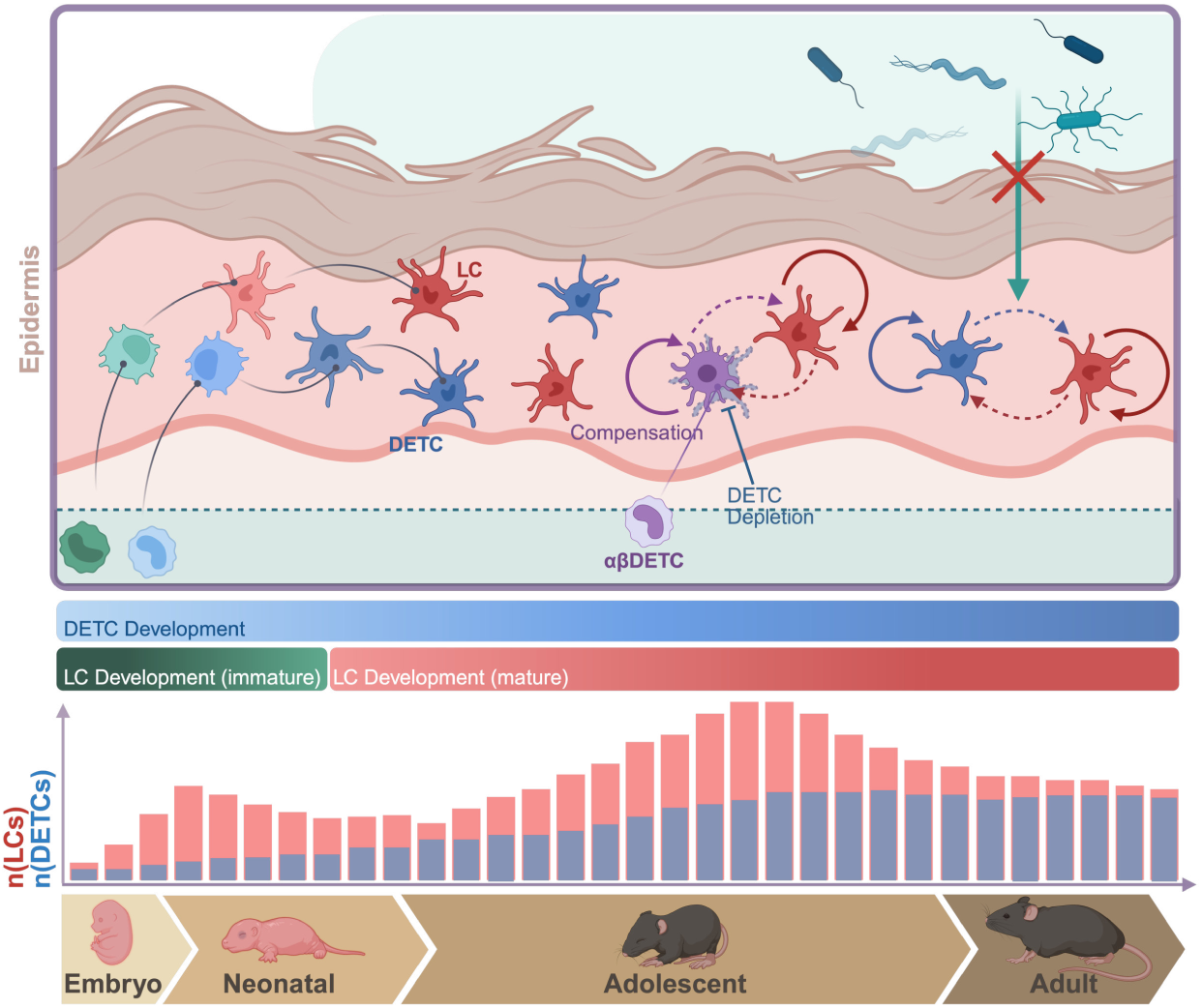

## INTRODUCTION

The skin is one of the largest and most complex organs in vertebrates. It separates the organism from the environment. It provides protection against dehydration, pathogens, toxins, and ultraviolet radiation, while it enables sensory perception of stimuli such as pain, heat, and touch at the same time (Segre, 2006). Beyond its structural role in maintaining hydration and shielding internal tissues, the skin functions as a highly specialized immune organ. Anatomically, the skin is divided into two major layers—the epidermis and the dermis—both of which host diverse immune cell repertoires that contribute to tissue homeostasis, immune surveillance, and tolerance to commensal microbiota (Belkaid & Tamoutounour, 2016; Pasparakis *et al*, 2014). The epidermis forms the outermost layer of the skin and consists of tightly packed, interconnected avascular layers of keratinocytes (Simpson *et al*, 2011). Despite its compact structure, the mouse epidermis contains a distinct and relatively stable resident immune cell community composed of two key populations: dendritic epidermal T cells (DETCs), unique γδ T cells expressing an invariant Vγ5Vδ1 T cell receptor (TCR) lacking junctional diversity (according to the Heilig and Tonegawa nomenclature) (Nielsen *et al*, 2017; Heilig & Tonegawa, 1986; Bergstresser *et al*, 1983), and Langerhans cells (LCs), a specialized subset of dendritic cell-like tissue-resident macrophages (Klareskog *et al*, 1977; Rowden *et al*, 1977; Stingl *et al*, 1977; Doebel *et al*, 2017).

In steady state, DETCs are critical for epidermal integrity, since their absence leads to increased keratinocyte apoptosis, likely due to reduced production of insulin-like growth factor I (IGF1) (Sharp *et al*, 2005). Moreover, LCs mediate tolerance to commensal microbiota, preemptive immunity, and immune surveillance (Seneschal *et al*, 2012; Ouchi *et al*, 2011). Their absence in adulthood has been associated with disorganized keratinocyte layering (Park *et al*, 2021), the emergence of autoreactive T cells (Kaplan *et al*, 2005), and compromised barrier function (Kubo *et al*, 2009). The functions of DETCs and LCs have been extensively studied in mouse models of skin inflammation, wound healing, and cancer. DETCs contribute to wound repair by sensing stressed keratinocytes and producing cytokines and growth factors such as interferon (IFN)-γ, tumor necrosis factor (TNF)-α, and keratinocyte growth factor (KGF) (Komori *et al*, 2012; Keyes *et al*, 2016; Yoshida *et al*, 2012; Jameson *et al*, 2002). LCs, in turn, act as antigen-presenting cells during infection and inflammation and are critical for preventing autoimmunity and limiting inflammatory responses (Kim *et al*, 2016; Kamenjarin *et al*, 2023). Dysregulation of either cell type in the adult epidermis has been associated with impaired tissue repair and inflammatory skin diseases (Jameson & Havran, 2007; Nielsen *et al*, 2017).

Both DETCs and LCs populate the mouse epidermis in the perinatal phase, entering the tissue niche in parallel or shortly after one another (Jiang *et al*, 2010; Gentek *et al*, 2018; Chorro *et al*, 2009; Chang-Rodriguez *et al*, 2005). Once established, they remain within this niche throughout life, maintaining their populations via endogenous proliferation in steady state (Kaplan, 2017; Chodaczek *et al*, 2012; Henneke *et al*, 2021; Ghigo *et al*, 2013). *Bona fide* adult LCs are thought to originate from erythromyeloid progenitors (EMPs) in the embryonic yolk sac. Between E8.5 and E10.5, EMPs colonize the fetal liver and give rise to fetal monocytes, which later migrate to the dermis. Around birth, LC progenitors transmigrate into the developing epidermis (Hoeffel *et al*, 2012; Chorro *et al*, 2009). DETCs, in contrast, derive from fetal thymic precursors, where they are the first T cells to develop in mouse. These precursors may themselves arise from the yolk sac (Gentek *et al*, 2018). Following epidermal entry, both DETCs and LCs undergo postnatal adaptation with the developing niche, acquiring their characteristic ramified morphology and functional properties (Chorro *et al*, 2009; Romani *et al*, 1985). Prior studies have identified essential signals instructing LC differentiation, including transforming growth factor-β (TGF-β1), and its receptor, interleukin-34 (IL-34), and colony stimulating factor 1 receptor (CSF1R) signaling (Borkowski *et al*, 1996; Chopin *et al*, 2013; Wang *et al*, 2012; Greter *et al*, 2012). While these pathways are indispensable for the generation and maintenance of adult LCs, Skint1 has been shown to be central for DETC development and differentiation in the mouse epidermis (Bauer *et al*, 2012; Lewis *et al*, 2006; Turchinovich & Hayday, 2011; Barbee *et al*, 2011; Boyden *et al*, 2008).

In the adult mouse skin, it was suggested that DETC and LCs contribute to coordinated spatial tiling and reciprocal signaling. DETCs can promote LC proliferation via cytokines, while LCs may influence DETC activation through antigen presentation and co-stimulatory signals (Park *et al*, 2021). However, the regulatory cues governing the postnatal differentiation of DETCs and LCs, as well as the longitudinal dynamics of this process, remain incompletely understood. In particular, it is still unclear whether their postnatal development is guided by shared or distinct signaling pathways and how these cell types interact in this critical period of life (Mohammed *et al*, 2016). Whether such crosstalk also contributes to their postnatal establishment and differentiation, however, remains largely unexplored.

Moreover, as both DETCs and LCs acquire immunophenotypic and functional properties around birth, it has been debated whether direct or distant (gut) colonization with commensal microbiota might influence this process, e.g. via microbial metabolites (Brand *et al*, 2023; Belkaid & Harrison, 2017; Henneke *et al*, 2021; Li *et al*, 2011). This has been well studied in the oral mucosa—where microbiota have been shown to shape gingival LC development and contribute to myeloid heterogeneity (Capucha *et al*, 2018; Jaber *et al*, 2023). In contrast, the impact of microbiota on epidermal LC and DETC development remains elusive. To date, however, no such studies have addressed the epidermal LCs. In contrast, prior studies have shown that DETC frequencies in the epidermis remain largely unaltered in germfree (GF) mice (Chodaczek *et al*, 2012; Papotto *et al*, 2021), suggesting that DETC development may be less dependent on microbial colonization. Thus, whether and how commensal microbes modulate the postnatal development of epidermal DETCs and LCs remains to be explored.

In this study, we present a comprehensive temporal map of DETCs and LCs in the mouse epidermis, spanning from late embryogenesis through postnatal development. We delineate distinct waves of population expansion, morphological remodeling, and transcriptional programming that shape these resident immune compartments postnatally. Our study provides a transcriptome-resolved single-cell atlas of postnatal DETC and LC development in the mouse epidermis, uncovering distinct and progressive developmental trajectories. Furthermore, we show that LC differentiation can proceed independently of canonical DETCs in the epidermis, most likely rescued by the presence of αβDETCs. Our analyses further revealed that the postnatal specification of both DETCs and LCs occurs independently of microbial cues. Taken together, our study highlights a parallel, but potentially independent development of the mouse epidermal immune compartment during early life.

## RESULTS

### Distinct Postnatal Expansion and Morphological Maturation of DETCs and LCs

To understand the patterning dynamics of DETCs and LCs across peri- and postnatal development, we performed high-resolution whole-mount imaging of mouse epidermis from embryonic day 17.5 (E17.5) to postnatal day 60 (P60) (Figure 1a). As previously demonstrated, LCs dynamically regulate marker expression during postnatal development. While they strongly express the chemokine receptor CX3CR1 in the early postnatal period, this expression declines over time, concurrent with the acquisition of differentiation markers such as epithelial cell adhesion molecule (EpCAM), Langerin (CD207), and MHC-II at varying rates (Suppl. Figure 1a, Figure 1b) (Chorro *et al*, 2009). In contrast, DETCs continuously express CX3CR1 and CD3ε throughout development (Havran & Allison, 1990; Almeida *et al*, 2015). Since both DETCs and LCs express CX3CR1 in the first postnatal week, we used the *Cx3cr1*^GFP^ reporter line in combination with CD3ε staining to discriminate both populations (Suppl. Figure 1a). Notably, LCs expressed low levels of Langerin around P4, which was subsequently used as a definitive marker for their identification and quantification (Suppl. Figure 1a). At E17.5, both DETCs and LCs were present in low numbers in the epidermis and exhibited rounded, immature morphologies (Figure 1b, c, Suppl. Figure 1a). By P0, LCs showed an expansion in cell number, while DETC numbers remained low (Figure 1b, c). Already at this early stage, LCs started forming initial protrusions (Figure 1b). Between P1 and P7, LCs further extended dendritic processes and exhibited increased ramification (Figure 1b, Suppl. Figure 1a). In contrast, DETCs developed their characteristic dendritic morphology more gradually, with prominent ramification observed only by P30 (Figure 1b, Suppl. Figure 1a). Quantification of cell numbers over time revealed that DETCs expanded steadily, reaching a plateau around P7, and maintained stable cell numbers in the epidermis into adulthood (Figure 1c). LCs, in contrast, displayed a more dynamic expansion pattern characterized by two waves: an initial increase around birth, a short plateau phase, followed by a pronounced proliferative burst at P7, and a subsequent decline until reaching adult equilibrium (Figure 1c). By P60, DETCs and LCs converged to comparable densities within the epidermis (Figure 1c).

**Figure 1.**
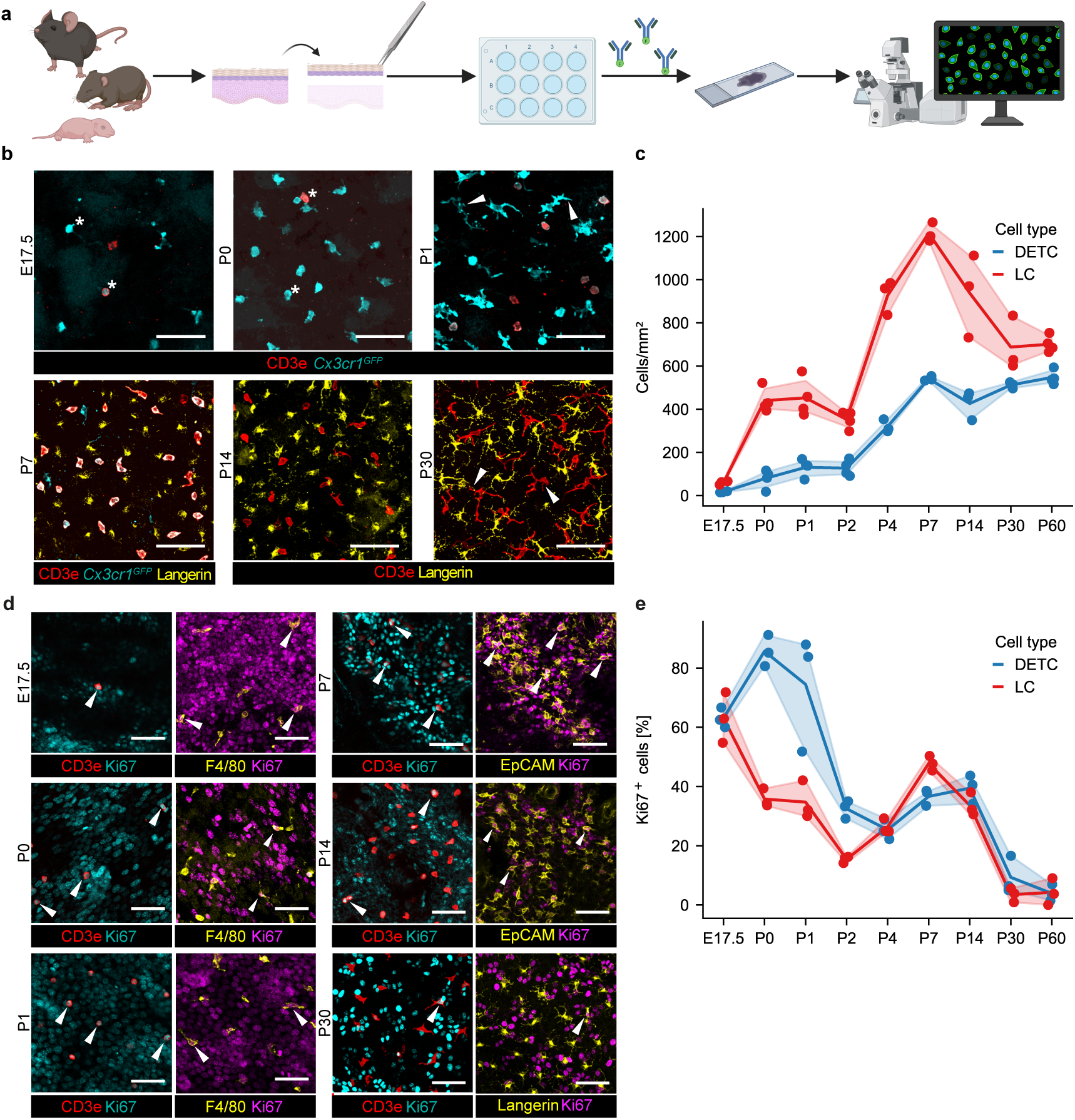
DETCs and LCs follow distinct expansion dynamics during postnatal development. **a** Schematic diagram of experimental flow for imaging analysis. **b** Representative whole mount confocal images of epidermal DETCs and LCs at different time points. CD3e is shown in red, *Cx3cr1^GFP^* is shown in cyan, and Langerin is shown in yellow. Asterisks indicate round cell states. Arrowheads indicate cell branching. Maximum projection of the confocal z-stack is shown. Scale bar= 50 µm. **c** Quantification of DETCs (blue) and LCs (red) across developmental stages. Each dot represents one animal. Results are representatives of three epidermal regions of one sample per mouse (n = 3 animals per group). **d** Representative whole mount confocal images of epidermal Ki67^+^ DETCs and LCs at different time points. CD3e is shown in red, Ki67 is shown in cyan or magenta, and F4/80 or Langerin are shown in yellow. Arrowheads indicate proliferating DETCs or LCs. Selected z layers are shown for illustration purposes. Scale bar= 50 µm. **e** Percentage of Ki67^+^ DETCs (blue) and LCs (red) across developmental stages. Each dot represents one animal. Results are representatives of three epidermal regions of one sample per mouse (n = 3 animals per group).

To assess the underlying proliferative dynamics of DETCs and LCs during this period, we labeled cycling cells (Ki67), combined with CD3ε for DETCs and F4/80 (E17.5–P4), EpCAM (P7–P14), or CD207 (P21–P60) for LCs (Suppl. Figure 1b). At E17.5, progenitors of both DETCs and LCs were highly proliferative, with ∼60% of cells in each compartment being Ki67⁺ (LCs: 63.17%±8.54% and DETCs: 63.06%±3.37%) (Figure 1d, e). By P0, LC proliferation rate declined (35.77%±3.13%), while nearly all DETCs remained Ki67⁺ (85.67%±5.29%), marking a proliferative peak for DETCs that persisted through the early postnatal days. At P7, LC proliferation rate increased again to (47.80%±2.47%), while Ki67⁺ DETCs decreased to (36.64%±2.76%), suggesting that DETCs were transitioning into a quiescent state, whereas LCs were entering their second proliferative wave (Figure 1d, e). In adulthood, only a small fraction of cells from either population remained Ki67⁺ (Figure 1d, e).

Together, these data show that DETCs and LCs undergo temporally distinct programs of proliferation, and morphological remodeling during postnatal development. Ramification, proliferation, and immune cell density are acquired by DETCs and LCs in a stepwise and differential manner, highlighting that each population follows a distinct differentiation trajectory within the developing epidermal niche.

### Temporal Single-Cell Transcriptomic Profiling of the Epidermal Immune Compartment

To gain deeper insights into the molecular postnatal programs governing DETCs and LCs, we performed single-cell RNA sequencing (scRNA-seq) of mouse epidermal immune cells across seven developmental stages, from embryonic day 18.5 (E18.5) to postnatal day 100 (P100) (Figure 2a). Building on our imaging-based characterization, we hypothesized that a transcriptomic approach would allow us to capture dynamic changes in gene expression and cell cycle status at single-cell resolution. We isolated single-cell suspensions from ventral epidermal sheets at early developmental stages (E18.5 to P7) and from ear epidermal sheets at later stages (P21 to P100). Following sample hashing to distinguish individual time points during analysis, CD45⁺ Gr1⁻ CD19⁻ single cells were FACS sorted and subjected to gene expression profiling using the 10x Genomics platform (Suppl. Figure 2a). After quality control and filtering (Suppl. Figure 2b), we obtained 51,737 high-quality cells across all stages (Figure 2b). Unsupervised clustering of transcriptomes revealed 18 transcriptionally distinct clusters that distributed across the full developmental timeline (Figure 2b, c, Suppl. Table 1).

**Figure 2.**
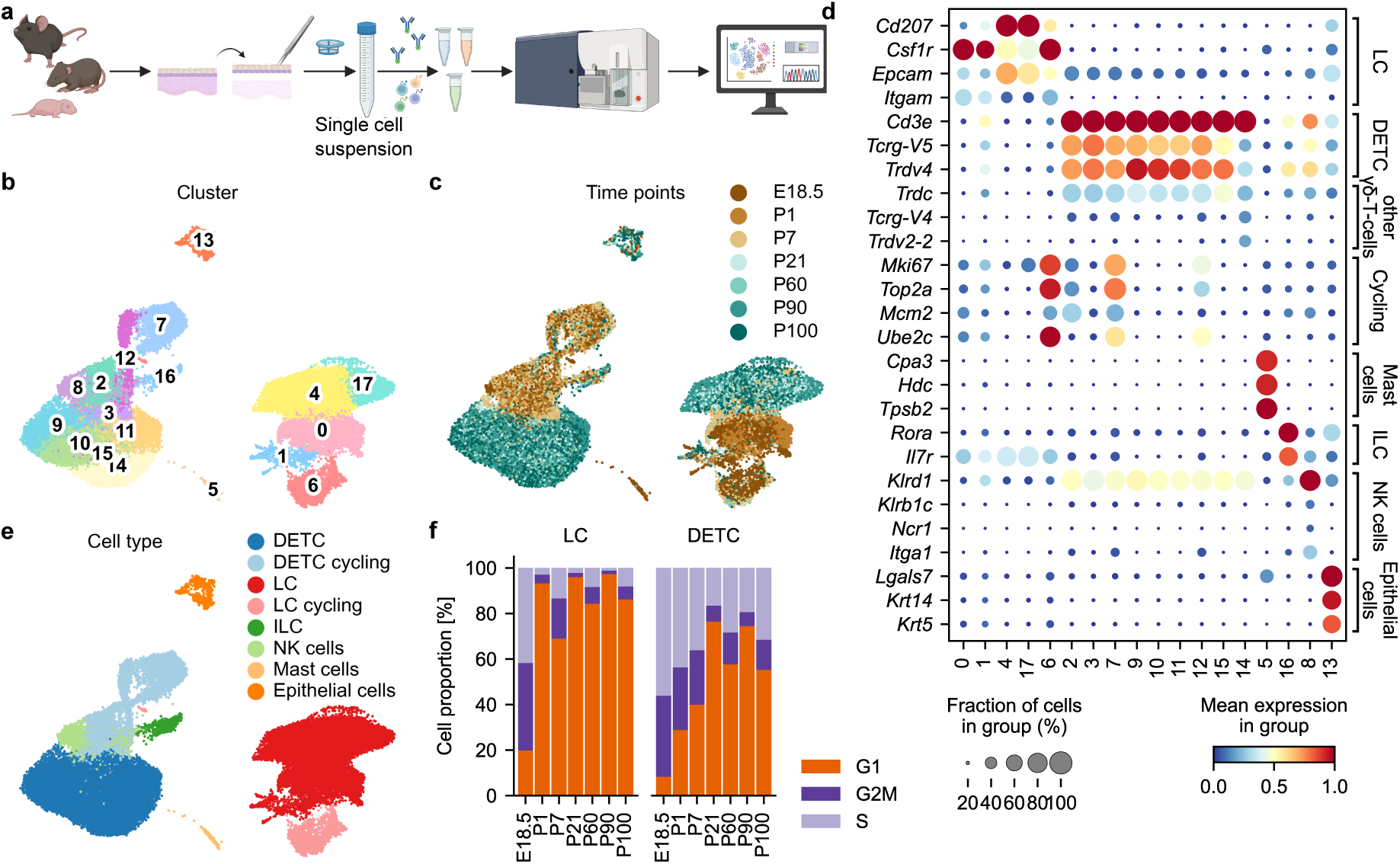
Single-cell transcriptomic profiling reveals distinct DETC and LC subsets during postnatal development. **a** Schematic diagram of experimental flow for scRNA-seq. **b** Uniform manifold approximation and projection (UMAP) plot of cell clusters after unsupervised clustering. Each color represents one cell cluster. (N = 51737 cells) **c** UMAP plot illustrating the distribution of single cells according to their time point of origin. **d** Dot plot showing key marker gene expression of cell types in the mouse epidermis across 18 identified clusters. Color represents the scaled mean expression of the gene in the respective cluster. Dot size represents the fraction of cells in the cluster expressing the gene. **e** UMAP plot displaying annotated immune cell types based on canonical marker gene expression. Each color represents one cell type. **f** Stacked bar plot illustrating the distribution of LCs (left) and DETCs (right) across G1 (orange), S (light purple), and G2/M (purple) cell cycle phases at each time point. The proportion of cells in each cell cycle phase is shown as a percentage of the total cell population.

Cell type annotation was performed using manually curated signature genes for skin-resident immune and non-immune populations (Figure 2d, e). Clusters 0, 1, 4, 6, and 17 expressed *Csf1r* and *Itgam* (encoding CD11b), consistent with a macrophage/LC identity. Among these, clusters 4 and 17 showed expression of LC markers *Cd207* and *Epcam*, which were largely absent in the other LC clusters (Figure 2d). Cluster 6 was enriched for cell cycle–associated genes such as *Mki67* and *Top2a*, indicating a highly proliferative subset within the LC compartment (Figure 2d). Clusters 2, 3, 7, 9, 10, 11, 12, 14, and 15 expressed *Cd3e* and *Trdc*, consistent with γδ T cell identity (Figure 2d). All clusters except cluster 14 expressed *Tcrg-V5* and *Trdv4*, identifying them as canonical Vγ5⁺Vδ1⁺ DETCs (Figure 2d). In contrast, cluster 14 expressed *Tcrg-V4* and *Trdv2-2*, marking them as Vγ4⁺ γδ T cells (Figure 2d). Additional immune populations were also detected. Cluster 5 expressed *Cpa3*, consistent with mast cells, whereas clusters 16 and 8 were annotated as innate lymphoid cells (ILCs) and natural killer (NK) cells, respectively, based on expression of *Rora*, *Il7r* (ILCs), and *Ncr1*, *Klrb1c* (NK cells). Cluster 13 showed high expression of *Epcam*, *Lgals7* (Pinto *et al*, 2023), *Krt14*, and *Krt5*, indicating minor epithelial/keratinocyte contamination (Figure 2d).

To characterize proliferative states during development, we performed cell cycle phase scoring using gene signatures specific for G1, S, and G2/M phases across all annotated DETCs and LCs (Figure 2f, Suppl. Table 2). Consistent with our imaging data, we observed a sharp transition from proliferative to quiescent states in both populations between birth and adulthood. Notably, LCs exhibited two distinct proliferative peaks: first at E18.5, with 38.4% of cells in G2/M-phase and 41.7% in S-phase, and a second at P7, with 17.7% of cells in G2/M and 13.3% in S-phase (Figure 2f). In contrast, DETCs showed a gradual decline in proliferative cells, with decreasing frequencies in both S and G2/M phases from E18.5 to P21 (Figure 2f). At later time points, both DETCs and LCs were predominantly in the G1 phase, consistent with entry into a quiescent state.

Together, these data confirm the presence of transcriptionally distinct DETC and LC subsets across development and reveal discrete waves of cell cycle activity. These findings reinforce the notion that DETCs and LCs follow temporally and molecularly distinct trajectories in the postnatal epidermis.

### DETCs Follow a Linear Maturation Trajectory in the Epidermis

Next, we focused on DETCs to characterize the transcriptional programs in postnatal development. Re-clustering of annotated DETCs within the scRNA-seq dataset identified eight transcriptionally distinct DETC clusters (Figure 3a). Clusters 0, 1, and 2 were enriched at early stages (E18.5, P1, P7), whereas clusters 3–7 were predominantly composed of cells from later time points (P21, P60, P90, P100), with cluster 3 being particularly overrepresented at P21 (Figure 3b, c). All clusters robustly expressed canonical DETC-associated genes, including *Il2rb* (CD122) and *Il2rg* (CD132), encoding IL-15 receptor subunits critical for DETC proliferation and survival (Edelbaum *et al*, 1995; Kawai *et al*, 1998), as well as *Thy1* (Bergstresser *et al*, 1983), *Xcl1* (Boismenu *et al*, 1996), and *Klrk1* (encoding NKG2D) (Whang *et al*, 2009) (Figure 3d, Suppl. Table 2). Key transcription factors involved in DETC development, including *Runx3* and *Ets1*, were also expressed across all clusters, albeit at lower levels (Woolf *et al*, 2007; Battaglia *et al*, 2023) (Figure 3d). Clusters 0, 1, and 2 showed high expression of cell cycle–associated genes such as *Mki67*, *Top2a*, *Mcm2*, and *Mcm6*, indicating highly proliferative states during early postnatal development, consistent with our imaging data (Figure 3d). In contrast, clusters 3–7, which were enriched in adult epidermis, expressed higher levels of T cell activation–related genes including *Nr4a2* and *Fos* (Figure 3d). Notably, DETCs upregulated the growth factor *Areg* in clusters 4–7—abundant after P21—suggesting the acquisition of tissue-repair functions during adulthood (Figure 3d) (du Halgouet *et al*, 2024). Moreover, clusters 4–7 also expressed genes with currently unknown roles in γδ T cell biology, such as *Ramp3*, *Gem*, and *Mest*, with cluster 5 showing the highest expression of all three. Next, we characterized distinct transcriptional programs in DETCs at different developmental stages using published gene sets (Figure 3e, Suppl. Table 3). This analysis revealed progressive upregulation of gene signatures related to structural and morphological remodeling, such as “dendrite development” (GO:0016358) and “cell projection organization” (GO:0030030), as well as tissue repair programs (Yanai *et al*, 2016; Arpaia *et al*, 2015; du Halgouet *et al*, 2023). Dendrite development and cell projection organization signatures increased gradually over time, whereas tissue repair programs were prominently induced from P60 onward, coinciding with adulthood.

**Figure 3.**
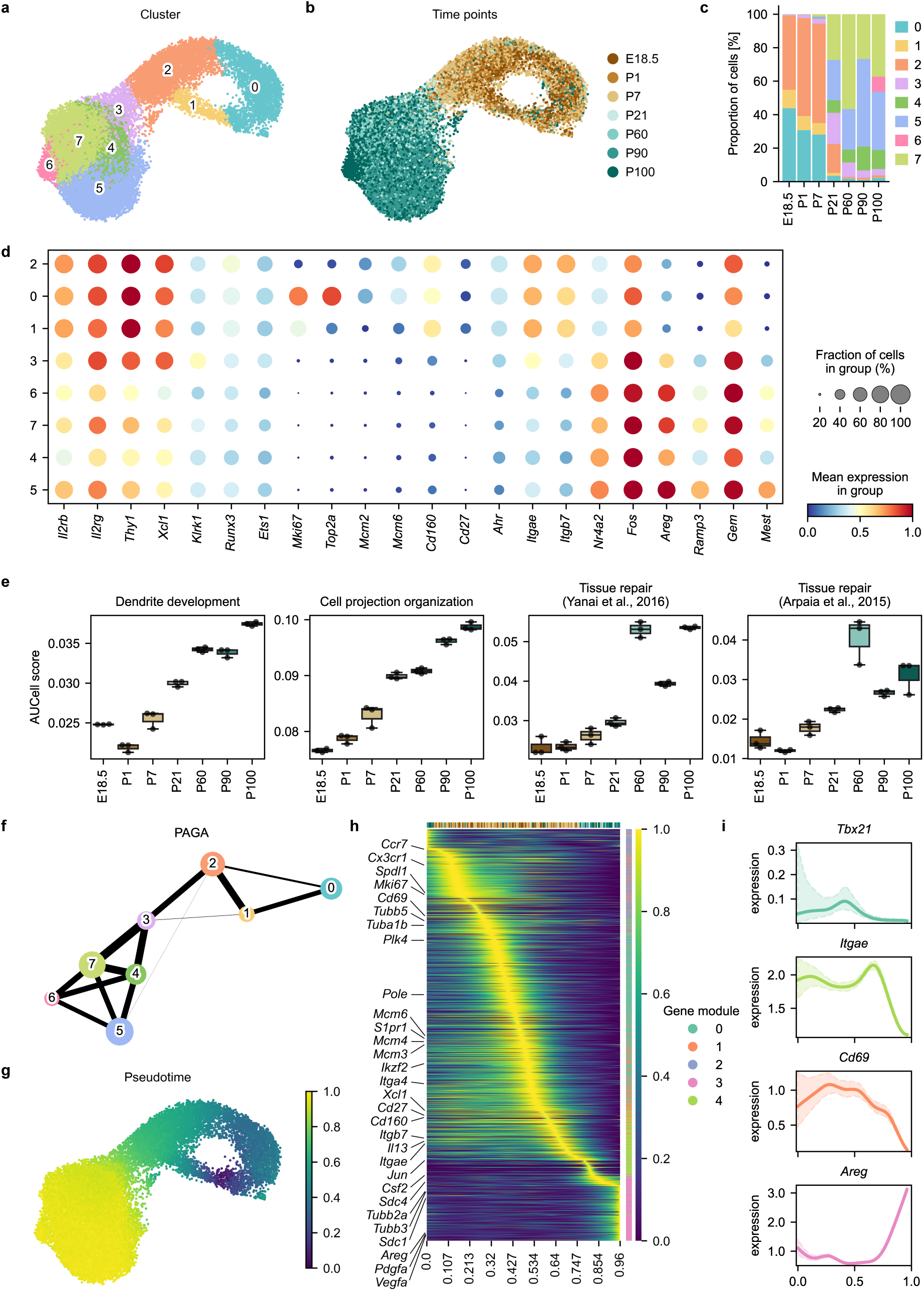
DETCs gradually acquire of tissue specificity and maturation markers in the developing epidermis. **a** UMAP plot of re-clustered DETCs. Eight clusters were identified (N = 25911 cells). Each color represents one cell cluster. **b** UMAP plot illustrating the distribution of cells according to their time point of origin. Each color represents one analysis time point. **C** Bar plot depicting the proportion of cells assigned to each cluster across the different time points. Colors indicate the respective clusters (see also Fig. 3a). **d** Dot plot showing expression of selected genes in the eight different DETC clusters. Color represents the scaled mean expression of the gene in the respective cluster and dot size represents the fraction of cells in the cluster expressing the gene. **e** Box plot showing mean AUCell score for GO terms and gene sets from previously published data (Yanai *et al*, 2016; Arpaia *et al*, 2015)) in DETCs across different time points. Mean +/- SEM is shown. Dots represent individual biological samples. **f** PAGA graph at cluster resolution. Each color represents one cell cluster (See also Fig. 3a). Each node represents one cluster, with node size proportional to the number of cells in the cluster. Edges indicate connectivity between clusters. Edge thickness reflects the strength of the connection based on transcriptomic similarities. **g** UMAP plot colored by diffusion pseudotime values calculated based on PAGA graph with the starting cell picked based on predicted initial state (Suppl. Figure 3a). Color scale reflects the relative progression of cells along developmental trajectories. Color scale is shown in legend. **h** Heatmap showing the smoothed expression of top 2000 highly variable genes ordered by their peak of expression along pseudotime (left to right) and organized into modules (top to bottom). Rows represent genes. Columns depict the pseudotime progression. Gene modules were identified based on expression patterns along the pseudotime. Five different gene modules were identified, and module membership is indicated by color in the vertical color bar. Second color scale depicts the relative progression of cells along developmental trajectories. Time points are indicated in the horizontal color bar above the heatmap according to the color code in **b**. Scale reflects the smoothed and scaled expression value of the genes. **i** Graphs showing the expression profile of individual, key genes along pseudotime from early to late stages (left to right). Colors indicate the respective module membership. Smoothed gene expressions are shown.

Next, we applied CellRank (Lange *et al*, 2022) to model DETC lineage progression based on our temporally annotated scRNA-seq dataset. We first constructed a real-time kernel using experimental time points to infer initial and terminal states of DETC differentiation (Suppl. Figure 3a, b). We then computed diffusion pseudotime (DPT) trajectories along a partition-based graph abstraction (PAGA) at the cluster level (Wolf *et al*, 2019) (Figure 3f), enabling modeling of gene expression changes over pseudotime (Figure 3g). To identify coordinated transcriptional programs, we smoothed gene expression along DPT and clustered the resulting expression trajectories into five distinct gene modules (Figure 3h, i, Suppl. Figure 3c, Suppl. Table 4). Gene modules 0 and 2, enriched at early to intermediate pseudotime, contained genes associated with S-phase and G2/M-phase of the cell cycle and DNA replication, including *Mcm3*, *Mcm4*, *Mcm6*, *Pole*, *Tuba1b*, *Tubb5*, *Mki67*, *Sdl1*, and *Plk4*. Gene module 1, expressed predominantly at early stages (until P7), was enriched for genes associated with IFN-γ–producing γδ T cell lineages, such as *Cd27*, and *Cd160*—possibly reflecting residual thymic programming (Figure 3h, i). This finding aligns with the enhanced IFN-γ production reported in neonatal DETCs compared to IL13-biased adult DETCs (Ibusuki *et al*, 2024). Module 1 also contained genes linked to T cell activation and migration, including *Cd69*, *Ccr7*, *Cx3cr1*, *Itga4*, and *S1pr1*, supporting the idea that DETCs retain migratory and activation signatures perinatally before transitioning to tissue residency (Figure 3h, i) (McKenzie *et al*, 2022). Gene module 4 showed a similar temporal pattern to module 1—high at early stages but declining more gradually (Suppl. Figure 3c). This module included genes that remained consistently expressed until adulthood, such as *Itgae*, *Itgb7*, *Csf2*, *Il13*, *Xcl1*, and *Jun*, suggesting roles in tissue anchoring, cytokine production, and baseline effector functions (Figure 3h, i). Finally, gene module 3, uniquely upregulated in differentiated DETCs, was enriched for genes associated with tissue repair (*Areg*, *Vegfa*, *Pdgfa*) and cytoskeletal remodeling (*Tubb3*, *Tubb2a*, *Sdc1*, *Sdc4*), potentially contributing to the acquisition of dendritic morphology and repair capacity during DETC maturation (Figure 3h, i).

Taken together, our analysis identifies a linear transcriptional trajectory for DETCs in the epidermis, characterized by a gradual loss of migratory and thymic features and the progressive acquisition of tissue-adaptive and repair-associated programs.

### LCs Undergo Stepwise and Programmed Differentiation in the Epidermis

To dissect the differentiation dynamics of LCs during postnatal development, we re-clustered all previously annotated LC single-cell transcriptomes (e.g., *Cd207*, *Csf1r*, *Epcam*, *Itgam*) (Figure 2b). Unbiased clustering revealed 11 transcriptionally distinct LC clusters across the developmental time course (Figure 4a, b). Clusters 0–5 were enriched in cells from embryonic and early postnatal stages (E18.5–P7), while clusters 6–10 were dominated by cells from later stages (P21–P100) (Figure 4b, c). At E18.5 and P1, clusters 0–2 were the most prevalent, but by P7 their frequencies markedly declined, coinciding with the emergence of later-stage clusters. Thus, P7 represents a major transcriptomic inflection point for LCs. To characterize these clusters, we identified differentially expressed genes (DEGs), revealing distinct transcriptional programs (Suppl. Figure 4a, b; Suppl. Table 2). Early-stage clusters (0–2 and 5) expressed high levels of macrophage progenitor genes, including *Cx3cr1*, *Adgre1*, *Fcgr1*, and *Mrc1* (Figure 4d). Notably, these clusters also showed expression of the microglial identity genes *Hexb*, *Apoe*, *Trem2*, *P2ry12*, and *Tmem119.* Thus early LC progenitors in the epidermis are transcriptionally reminiscent of fetal microglia-like progenitors recently reported in human epidermis, testis, and heart (Wang *et al*, 2023) (Figure 4d). These clusters also expressed key transcription factors such as *Id2*, *Spi1*, *Irf8*, and *Cebpb*. While *Id2*, *Spi1*, and *Irf8* promote LC differentiation (Schiavoni *et al*, 2004; Chopin *et al*, 2013), *Cebpb* has been implicated as a negative regulator (Iwama *et al*, 2002; Hashimoto-Hill *et al*, 2018). Complement system genes were prominently expressed in early clusters and rapidly declined by P7, suggesting roles in perinatal LC function. Core LC-maintenance genes such as *Csf1r* and *Tgfbr1* were expressed in all clusters but were elevated in early stages (clusters 0–5), accompanied by *Tgfb1* expression (Figure 4d). In contrast, *Il34* was undetectable and *Csf1* was transiently expressed only in early LC progenitors, indicating that these niche factors are likely provided by other skin-resident cells (Suppl. Figure 4c, d). The expression of *Csf2ra* and *Csf2rb* across all clusters suggests that LCs remain poised to respond to colony stimulating factor 2 (CSF2) (also called: granulocyte-macrophage colony stimulating factor (GM-CSF)) (Figure 4d). Clusters 3, 4, and 6–10, enriched in later developmental stages, expressed hallmark LCmarkers, including MHC-II related genes (e.g., *H2-Aa*, *H2-Eb1*), *Cd207*, and *Epcam* (Figure 4d).

**Figure 4.**
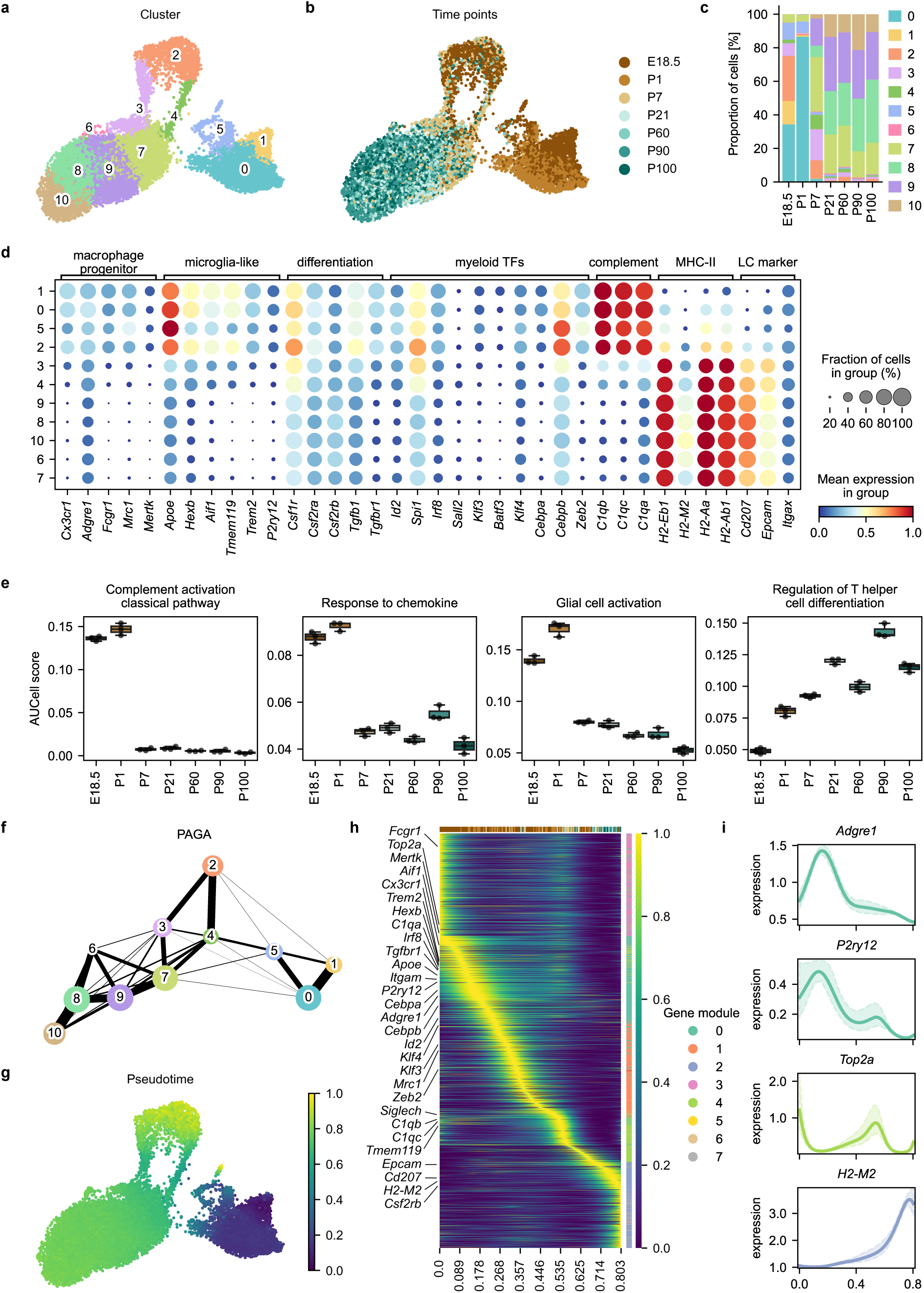
Stepwise maturation of LCs in the developing epidermis. **a** UMAP plot of re-clustered LCs. 11 clusters were identified (N = 17740 cells). Each color represents one cell cluster**. b** UMAP plot illustrating the distribution of cells according to their time point of origin. Each color represents one analysis time point. **c** Bar plot depicting the proportion of cells assigned to each cluster across the different time points. Colors indicate the respective clusters (see also Fig. 4a). **d** Dot plot showing expression of selected genes across the 11 different LC clusters. Color represents the scaled mean expression of the gene in the respective cluster and dot size represents the fraction of cells in the cluster expressing the gene. **e** Box plot showing mean AUCell scores per replicate of selected GO terms across the different time points. Mean +/- SEM is shown. Dots represent individual biological samples. **f** PAGA graph at cluster resolution. Each color represents one cell cluster (see also Fig. 4a). Each node represents one cluster, with node size proportional to the number of cells in the cluster. Edges indicate connectivity between clusters. Edge thickness reflects the strength of the connection based on transcriptomic similarities. **g** UMAP plot colored by diffusion pseudotime values calculated based on PAGA graph using predicted initial state as starting cell (Suppl. Figure 3e). Scale reflects the relative progression of cells along developmental trajectories. Color scale is shown in legend**. h** Heatmap showing the smoothed expression of top 2000 highly variable genes ordered by their peak of expression along pseudotime (left to right) and organized into modules (top to bottom). Rows represent genes. Columns depict pseudotime progression. Gene modules were identified based on expression patterns along the pseudotime. Eight different modules were identified, and module membership is indicated by color in the vertical color bar. Second color scale depicts the relative progression of cells along developmental trajectories. Time points are indicated in the horizontal color bar above the heatmap according to the color code in **b**. Scale reflects the smoothed and scaled expression value of the genes. **i** Graphs showing the expression profile of individual, key genes along pseudotime from early to late stages (left to right). Colors indicate the respective module membership. Smoothed gene expressions are shown.

To better understand functional transitions across development, we performed GO term enrichment analyses (Suppl. Table 5). At E18.5 and P1, early LCs were enriched for terms such as “complement activation classical pathway” (GO:0006958), “response to chemokine” (GO:1990868), and “leukocyte activation involved in inflammatory response” (GO:0002269) (Figure 4e, Suppl. Figure 4b, Suppl. Table 3). These signatures, alongside enrichment for “glial activation” (GO:0061900), reinforce the activated and microglia-like nature of LC progenitors. GO terms related to DNA replication (“DNA replication initiation”, GO:0006270) were also enriched at E18.5 and P7, consistent with our imaging data showing proliferative bursts at these time points (Figure 1d, e, Suppl. Figure 4b, Suppl. Table 3). Interestingly, genes linked to “regulation of T helper cell differentiation” were upregulated from P1 onwards, suggesting a potential role for LCs in influencing DETC polarization in the early epidermis (Figure 4e). In contrast, later stages were enriched for terms like “cell-cell junction assembly” (GO:0007043), indicating increasing involvement in structural organization and barrier function (Suppl. Figure 4b). Together, these data reveal a stepwise LC differentiation process during postnatal development, transitioning from a progenitor and microglia-like transcriptional program before P7 to a mature, tissue-integrated LC identity after P21. This is accompanied by two proliferative bursts—after epidermal colonization (E18.5) and at P7. Pseudotime analysis using PAGA and DPT, with cluster 1 at E18.5 as the root (Suppl. Figure 4e) revealed a bifurcated trajectory: one branch toward cluster 10 (dominant at later stages) and another toward proliferative cluster 2 (Figure 4f, g). Cluster 2 thus represents a key branching point that diverts cells from differentiation into a proliferative loop, which ultimately feeds into fully differentiated clusters (7–10). Our model does not exclude the possibility that mature LCs re-enter the cycle via this proliferative route, supporting the idea of intermittent expansion during LC differentiation. We further defined gene modules across pseudotime (Suppl. Figure 4f; Suppl. Table 4). Modules 0 and 3, active early, contained macrophage and microglial identity genes (e.g., *Fcgr1*, *Adgre1*, *P2ry12*, *Hexb*, *Tmem119*) (Figure 4h, i). Module 4, which peaked at early and intermediate stages, comprised proliferation-related genes such as *Top2a*. Module 2, enriched at the terminal trajectory, contained canonical LC markers (*Cd207*, *H2-M2*, *Epcam*), confirming the acquisition of a mature LC identity over time.

Collectively, our results support a model in which perinatal LCs development comprises discrete transcriptional states marked by successive functional modules—beginning with a macrophage progenitor identity, with a microglia-like signature, and culminating in full LC differentiation. This trajectory is interspersed with temporally regulated proliferative bursts.

### LC Development Proceeds Independently of Canonical DETCs

To assess whether DETCs influence LC development in the postnatal epidermis, we employed a computational cell–cell communication inference strategy using LIANA and Tensor-cell2cell (Dimitrov *et al*, 2024). Analyses were performed independently for each biological replicate across developmental time points, integrating predictions from multiple ligand–receptor inference methods into a unified consensus score. These were compiled into a four-dimensional tensor (sender cell, receiver cell, ligand–receptor pair, developmental context), followed by tensor decomposition, which revealed seven communication factors (Suppl. Figure 5a). Among these, factors 1, 5, 6, and 7 reflected prominent DETC–LC interactions across development (Figure 5a, Suppl. Table 6). While factors 5 and 7 represented predicted signals enriched in early development, factors 1 and 6 were predominant at later stages (Figure 5a). Notably, factors 1 and 6 included predicted LC-to-DETC signals at both early and late time points (Figure 5a, b). In addition to adhesion-related interactions, a strong prediction for *Ccl4*–*Ccr8* signaling between LCs and DETCs was found during early development (Figure 5b). However, CCL4 is not known to functionally bind CCR8, questioning the relevance of this interaction. Conversely, factors 6 and 7 captured DETC-derived signals to LCs, including cytokine–receptor pairs such as *Il13*→*Il13ra*_*Il4ra*, *Csf2*→*Csf2ra*_*Csf2rb*, and *Tgfb1*→*Tgfbr1*_*Tgfbr2*. These pathways are well-established in LC biology but have not been previously attributed to DETCs during development (Zhang *et al*, 2016, 2017; Kel *et al*, 2010; Caux *et al*, 1992; Witmer-Pack *et al*, 1987; Zheng *et al*, 2009) (Figure 5b). The *Tgfb1*→*Tgfbr1*_*Tgfbr2* interaction scored highly during early stages and waned over time. These results suggest DETCs may provide developmental cues for LC development in the early epidermis.

**Figure 5.**
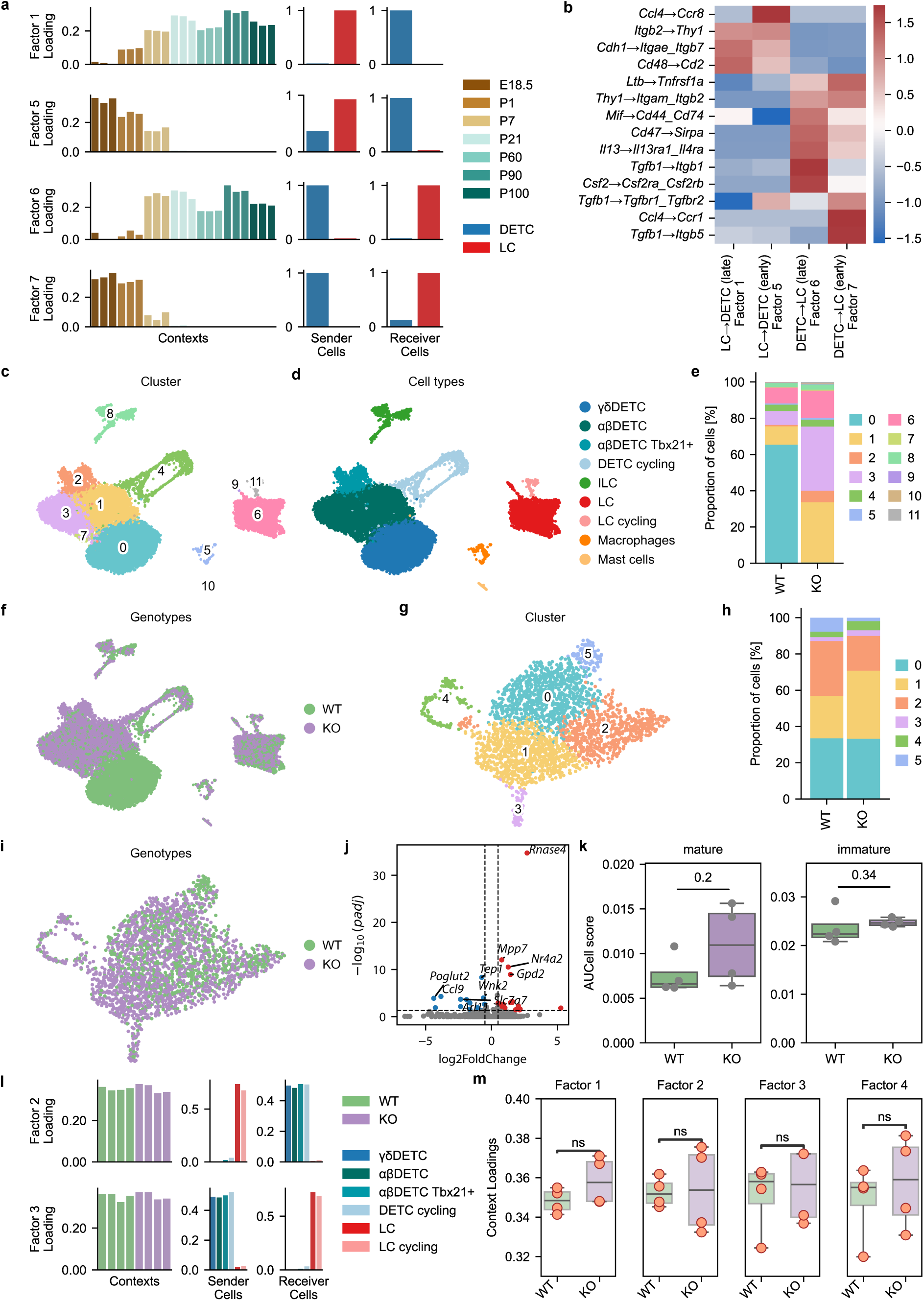
Loss of canonical DETCs during postnatal development does not affect the transcriptomic profile of adult LCs. **a** Graphs depicting results of running LIANA and Tensor-cell2cell frameworks. Each row represents a factor displaying paracrine signaling between DETCs (blue) and LCs (red), and each column a tensor dimension, wherein each bar plot represents an element of that dimension (time point, a sender cell, or a receiver cell). Factor loadings (y-axis) are displayed for each element of a given dimension. Colors of individual time points are depicted in legend. **b** Heatmap showing the loadings of selected interactions in factor 1, 5 -7 (paracrine interactions). Scale represents the loadings for an interaction in the factor. **c** UMAP plot of cell clusters after unsupervised clustering in *Tcrd^-/-^* (KO) (n = 4) and *Tcrd^+/+^* (WT) (n = 4) animals. 12 clusters were identified (N = 20632 cells). Each color represents one cell cluster. **d** UMAP plot showing different annotated cell types and cycling cells based on core marker expression. Cell type is indicated by color. **e** Bar plot showing the proportion of cells assigned to each cluster in KO and WT animals. Colors indicate the respective clusters (see also Fig. 5c). **f** UMAP plot showing cells colored by genotype of origin. Cells from KO mice are shown in purple, cells from WT mice are shown in green. **g** UMAP plot of re-clustered LCs. Six clusters were identified (n = 2606 cells). Each color represents one cluster. **h** Bar plot showing the proportion of cells assigned to each cluster originating from either KO or WT animals. Colors indicate the respective clusters (see also Fig. 5g). **i** UMAP plot showing cells colored by genotype. Cells from KO mice are shown in purple, cells from WT mice are shown in green. **j** Volcano plot of differentially expressed genes in LCs from KO vs WT mice as determined using PyDESeq2 (pseudobulked gene expression). Vertical lines mark a log2 fold change of 0.5 (fold change of ∼±1.41) and horizontal line represents a -log10 p-value of 1.3 (p-value of 0.05). Genes in red are significantly upregulated in LCs of KO mice and genes in blue are significantly downregulated. **k** Box plots of mean AUCell scores of gene sets for mature and immature LCs (signatures derived from Fig. 4) in WT and KO mice (n = 4). Mean +/- SEM is shown. Dots represent individual biological replicates. Statistical testing was performed using Mann-Whitney U rank test. **l** Graphs depicting results from running Tensor-cell2cell frameworks using the αβDETCs and γδDETCs and LCs from KO (purple) and WT mice (green). Each row represents a factor displaying paracrine signaling, and each column a tensor dimension, wherein each bar plot represents an element of that dimension (genotype, sender or receiver cells). Factor loadings (y-axis) are displayed for each element of a given dimension. Legend depicts color code for genotypes and cell types. **m** Box plots of context loading separated by genotype KO (purple) vs. WT (green) for all four factors derived from tensor decomposition. Dots represent individual biological samples. Mean +/- SEM is shown. Statistical testing was performed using Mann-Whitney U rank test with Benjamini-Hochberg correction for multiple testing.

To test whether DETCs are functionally required for LC development, we analyzed *Tcrd*-deficient (*Tcrd*^⁻/⁻^) mice, which lack all γδ T cells (Itohara *et al*, 1993). In these mice, DETCs are replaced by compensatory αβ T cells (hereafter αβDETCs) (Binz *et al*, 2021). We performed scRNA-seq on epidermal immune cells from adult *Tcrd*^⁺/⁺^ and *Tcrd*^⁻/⁻^ mice. After quality control, DETCs and LCs were identified in both genotypes, with small subsets of cycling cells within each population (Figure 5c, d, Suppl. Table 1). As expected, *Tcrd*^⁻/⁻^ mice lacked γδDETCs (cluster 0) but harbored abundant αβDETCs (cluster 1), whereas ILCs, mast cells, and macrophages were equally present in both genotypes (Figure 5c–f). Next, we sub-clustered LCs and identified 11 transcriptionally distinct clusters that were shared across both genotypes without genotype-specific alterations (Figure 5g–i). Pseudobulk differential expression analysis using pyDESeq2 (accounting for batch effects) revealed only 66 DEGs between *Tcrd*⁺/⁺ and *Tcrd*⁻/⁻ LCs, with modest fold changes (Figure 5j, Suppl. Table 1). While some of these genes were associated with LC or DC maturation, they did not indicate a substantial disruption of developmental programs. To quantify LC maturation, we constructed gene signatures for immature and mature LCs based on our time-course dataset and computed maturity scores. These scores did not differ significantly between genotypes (Figure 5k), suggesting that γδDETCs are not essential for LC maturation. To further explore whether αβDETCs functionally compensated for absent γδDETCs, we re-applied LIANA and Tensor-cell2cell using genotype as the contextual variable and modeled interactions among γδDETCs, αβDETCs, and LCs. Four communication factors were identified, yet none displayed genotype-specific signaling patterns (Figure 5l, m; Suppl. Figure 5c, Suppl. Table 6). Notably, predicted interactions such as *Il13*→*Il13ra*_*Il4ra*, *Csf2*→*Csf2ra*_*Csf2rb*, and *Tgfb1*→*Tgfbr1*_*Tgfbr2* were shared between γδDETCs and αβDETCs, with comparable interaction scores across genotypes (Figure 5m).

Together, these findings indicate that DETCs may contribute to a cytokine-rich microenvironment conducive to LC development. However, αβDETCs can recapitulate key signaling features of γδDETCs and may support LC differentiation.

### Commensal Microbiota Are Dispensable for DETC and LC Maturation

As the skin is among the first organs to be colonized by commensal microorganisms (Cha *et al*, 2025), we next asked whether the microbiota influenced the postnatal maturation of DETCs and LCs. To address this, we studied GF mice, which are devoid of microbial colonization, and wildlings, natural microbiota-based mouse model, which harbors a complex and naturally co- evolved microbiota and fungi resembling that of wild mouse counterparts (Rosshart *et al*, 2019; Runge *et al*, 2025; Bruno *et al*, 2025). We profiled epidermal immune cells at early (P1, P7) and adult (P90, P100) stages in specific-pathogen free (SPF), GF, and wildlings to assess both immediate and long-term microbiota-driven effects (Figure 6a, b, Suppl. Figure 6a, b). DETCs and LCs were annotated based on previously defined gene signatures from our time-course dataset (Figure 2, Suppl. Table 1). Immature DETCs were identified by *Cd160*, *Itgae*, and *Itgb7*, and mature DETCs by *Areg*, *Ramp3*, and *Mest*. Immature LCs were marked by complement-associated genes (*C1qb*, *C1qc*) and microglia-related transcripts (*Trem2*), while mature LCs expressed *Cd207*, *Epcam*, and MHC-II genes including *H2-Eb1*. Cycling DETCs and LCs were defined by proliferation-associated genes such as *Mki67* and *Top2a* (Figure 6c, Suppl. Figure 6c, d).

**Figure 6.**
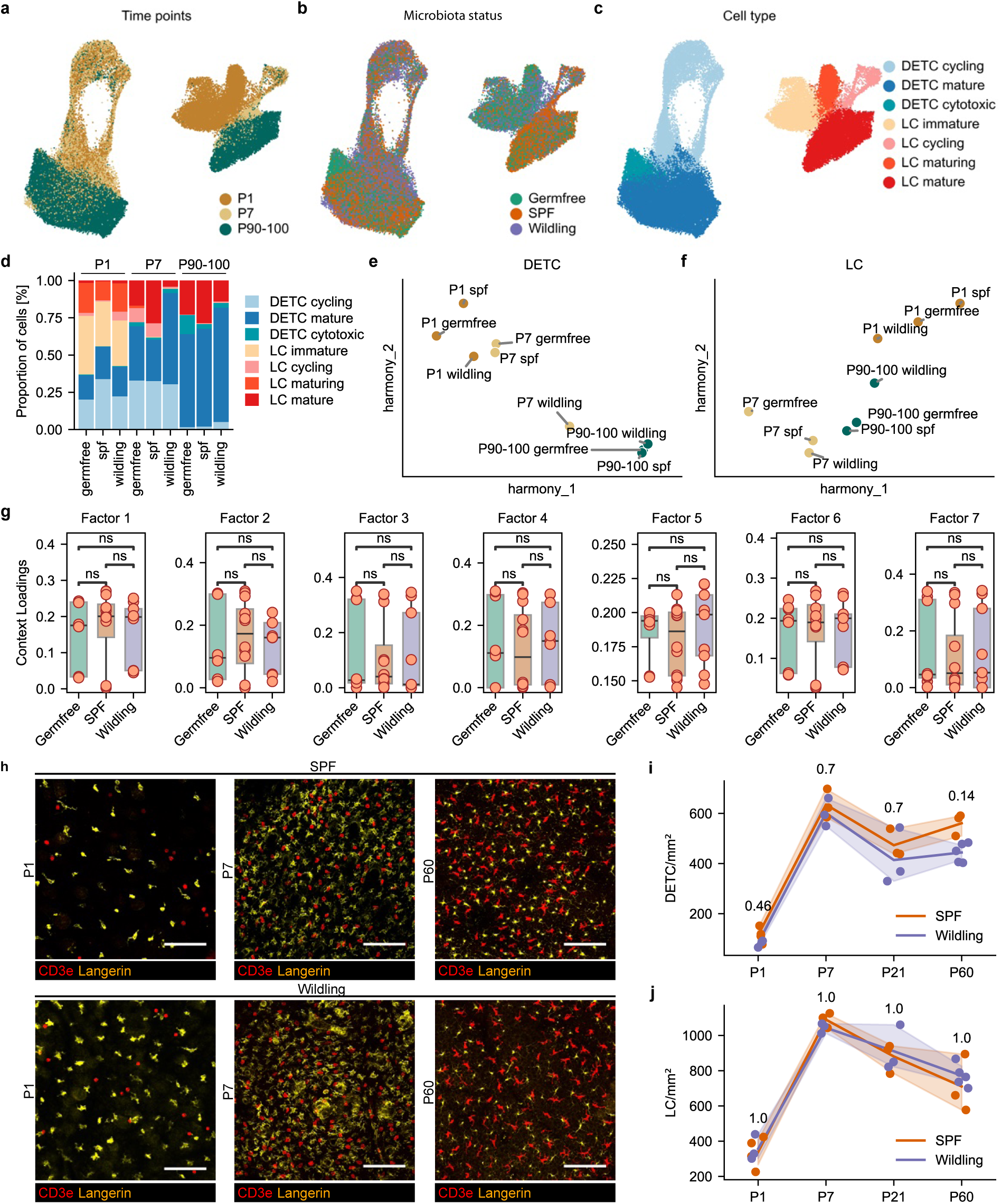
Absence or higher complexity of commensal microbiota does not affect DETC and LC maturation. **a** UMAP plot illustrating the distribution of cells from germ-free (GF) mice (n = 3), SPF mice (n = 3) and wildlings (n = 3) (N = 80556 cells) according to their time point of origin. Each color represents one analysis time point. **b** UMAP plot depicting the distribution of cells according to their microbiota status (GF (green), SPF (orange), wildlings (purple)). **c** UMAP plot showing different annotated DETCs and LCs, indicating cycling and differentiation states based on core marker expression (based on Suppl. Fig. 6d). Cell type is indicated by color. **d** Bar plot showing the proportion of cells within the different defined states (annotated in Fig. 6c) across different microbiota statuses (GF, SPF, wildlings) and analyzed time points. Time points are indicated above bar plot. Colors represent cell types (see also Fig. 6c). **e** Scatter plot showing the median of the harmony embedding for each time point and microbial status for DETCs. Colors indicates time points. Microbial status is labeled within the scatter plot. **f** Scatter plots showing the median of the harmony embedding for each time point and microbial status for LCs. Colors indicates time points. Microbial status is labeled within the scatter plot. **g** Box plots of the context loadings for all seven factors recovered by factor decomposition of a tensor built using microbial status and time points as contexts. Mean +/-SEM is shown. Statistical testing was performed using Mann-Whitney U rank test with Benjamini-Hochberg correction for multiple testing. **h** Representative images of epidermal DETCs and LCs in SPF and wildlings at P1, P7 and P60. Langerin is shown in orange and CD3e is shown in red. Maximum projection of the confocal z-stack is shown. Scale bar= 50 µm. **i** Quantification of DETC densities in SPF (orange) and wildling mice (purple) across developmental stages. Each dot represents one animal. Results are representative of three epidermal regions of one sample per mouse (n = 3 animals per group). Statistical significance was assessed using the Mann–Whitney U test, with p-values corrected for multiple testing using the Benjamini–Hochberg procedure. **j** Quantification of LC densities in SPF (orange) and wildlings (purple) across developmental stages. Each dot represents one animal. Results are representative of three epidermal regions of one sample per mouse (n = 3 animals per group). Statistical significance was assessed using the Mann–Whitney U test, with p-values corrected for multiple testing using the Benjamini–Hochberg procedure.

At P1, both DETCs and LCs displayed similar transcriptomic profiles across SPF, GF, and wildlings (Figure 6d). At P7, wildlings showed a modest reduction cell clusters associated with differentiated LCs and a corresponding increase in mature DETCs; however, this effect was no longer evident by P90–P100 (Figure 6d). Harmony-based clustering confirmed that developmental time point, and not microbial status, was the primary driver of transcriptional variation in both cell types (Figure 6e, f). To further explore potential microbiota-driven alterations in cell–cell communication, we again applied the LIANA and Tensor-cell2cell frameworks. Tensor decomposition revealed seven communication factors (Suppl. Figure 7a, Suppl. Table 6), none of which differed significantly across microbial conditions (GF, SPF, wildlings) (Figure 6g). Instead, observed variation in communication patterns was entirely attributable to developmental stage (Suppl. Figure 7a, b). To validate these transcriptomic findings, we performed immunofluorescent staining of DETCs and LCs across developmental time points in SPF mice and wildlings. In line with previous reports showing comparable mucosal DETC and LC frequencies in GF and SPF animals (Capucha *et al*, 2018; Chodaczek *et al*, 2012; Papotto *et al*, 2021), we found no significant differences between SPF mice and wildlings (Figure 6h-j).

Taken together, these results indicate that although microbial colonization may exert transient effects on LC maturation during early postnatal life, both DETCs and LCs ultimately mature independently of the microbiota in the epidermis. Thus, commensal microbes are dispensable for their postnatal development under steady-state conditions.

## Discussion

Our study provides a comprehensive temporal roadmap of the perinatal and postnatal development of DETCs and LCs in the mouse epidermis. By integrating high-resolution whole-mount imaging, and single-cell transcriptomics, we delineate distinct developmental trajectories of these resident immune populations, characterized by successive waves of proliferation, morphological remodeling, and transcriptional programming. Our findings demonstrate that LC maturation can proceed independently of canonical DETCs and that both cell types develop independent of commensal microbial colonization, suggesting a robust perinatal immune development program.

Our results align with previous studies demonstrating that DETC and LC progenitors colonize the mouse epidermis during the perinatal period (Gentek *et al*, 2018; Hoeffel *et al*, 2012). We detected both progenitor populations in the developing epidermis around E17.5. Thus, colonization by one cell type is not required for robust epidermal entry of the other, consistent with prior reports showing LC colonization within the same developmental window in Rag^−/−^ γc^−/−^ mice lacking DETCs (Chorro *et al*, 2009). However, we observed distinct postnatal expansion dynamics for DETCs and LCs. DETCs gradually increased in density until reaching their adult network configuration, whereas LCs underwent two proliferative waves, at E18.5 and P7. LC numbers peaked at P7 and subsequently declined until adulthood – a pattern also observed during postnatal development in other tissue-resident macrophage populations such as microglia in the CNS parenchyma (Barry-Carroll *et al*, 2023; Masuda *et al*, 2022). The mechanisms underlying this initial expansion and subsequent contraction of the population size remains to be defined. Interestingly, we also observed differences in the kinetics of dendrite formation: LCs began branching earlier than DETCs, suggesting that LCs may initiate tissue surveillance programs earlier during development.

Similar to the distinct expansion dynamics observed for both cell types, we found that DETCs underwent a gradual, tissue-specific maturation in the postnatal epidermis, whereas LCs progressed through a stepwise differentiation process, engaging distinct genetic programs. Early-stage DETCs exhibited high expression of cell cycle–associated and migratory genes, as well as markers of IFN-γ–producing γδ T cell lineages, consistent with residual thymic imprinting and a cytotoxic bias. This is in line with previous studies showing that neonatal DETCs preferentially produce IFN-γ upon activation in the perinatal epidermis (Ibusuki *et al*, 2024; Dalessandri *et al*, 2016). During DETC maturation, they progressively upregulate genes involved in cytoskeletal organization and tissue repair, along with increased expression of gene sets related to dendrite development—reflecting their adaptation to a homeostatic surveillance role. In addition to known maturation programs, we also identified genes with poorly characterized roles in DETCs, such as *Ramp3*, *Gem*, and *Mest*. *Ramp3*, encoding a receptor activity–modifying protein that regulates calcitonin gene–related peptide (CGRP) receptor trafficking, which has been linked to neuropeptide signaling in T cells, where it promotes T helper 1 (Th1) differentiation (Hou *et al*, 2024). *Gem* (GTP-binding protein expressed in mitogen-stimulated T cells) is induced upon T cell activation and influences cytoskeletal remodeling, chemotaxis, and actin dynamics (Chevalier *et al*, 2014; Maguire *et al*, 1994). In contrast, *Mest* (mesoderm-specific transcript), an imprinted gene associated with adipogenesis and mesenchymal development, has no established role in immune cells but may regulate epigenetic or metabolic programs (Kaneko-Ishino *et al*, 1995; Takahashi *et al*, 2005; Nikonova *et al*, 2008). The enrichment of these genes in mature DETCs suggests potential contributions to niche adaptation or immunometabolic homeostasis, although their precise functions in γδ T cells remain to be elucidated.

In contrast to our observations in DETCs, we identified distinct gene programs in LCs across development, suggesting a stepwise differentiation process marked by gene profile switches. First, our analyses demonstrated that LCs at E18.5 and P1 share many characteristics with activated macrophages and immature macrophage progenitors. Early LC progenitors are derived from fetal monocytes, migrate to the dermis at E16.5, and differentiate into macrophage progenitors within the tissue before entering the epidermis and developing into LCs (Chorro *et al*, 2009; Hoeffel *et al*, 2012). Interestingly, we found that LC progenitors transiently express core microglial signature genes and gene programs during the perinatal period, which disappear shortly thereafter. Recent studies have reported microglia-like progenitors in human fetal skin, testis, and heart (Bian *et al*, 2020; Gopee *et al*, 2024; Wang *et al*, 2023). These microglia-like progenitors display signatures reminiscent of the microglial programs we observed in mouse LCs during the perinatal window. This suggests a specific imprinting of recruited progenitors by epidermal niche factors such as IL-34 or TGF-β1, analogous to imprinting mechanisms in the developing central nervous system. However, despite the continuous presence of these niche factors throughout postnatal development, the microglia-like signature in LCs vanishes shortly after birth. (Wang *et al*, 2016; Kaplan *et al*, 2007). The stepwise developmental diversion of pre- and perinatally seeded skin immune cells away from their fetal programs is in line with that of dermal macrophages that mostly loose CX3CR1 expression with enduring tissue residency (Kolter *et al*, 2025). The transient nature of this microglial signature may indicate that LC progenitors interact with neurons during early epidermal colonization—for example, by engaging with outgrowing sensory neurons in the dermis or utilizing them as migratory tracks to access the epidermis (Gopee *et al*, 2024). This is supported by previous reports of sciatic nerve–associated macrophages exhibiting microglia-like gene signatures, despite eventually acquiring a niche-specific identity (Ydens *et al*, 2020). Thus, dermal sensory neurons which harbor sensory neuron–associated macrophages may also serve as an anatomical hub for LC progenitors prior to their migration into the epidermis (Kolter *et al*, 2025, 2019). Our data further indicate that core programs associated with LC maturation are established from P7 onward, as evidenced by the upregulation of known LC marker genes such as *Cd207*, *Epcam*, and MHC-II–related genes. This supports the idea of early tissue imprinting that culminates around the time of weaning (P21), which appears to represent a developmental checkpoint for LC maturation. Subsequently, the acquisition of tissue-specific functions—such as cell–cell junction formation, reinforcement of epidermal barrier integrity, and immune surveillance—is further associated with the completion of this maturation process.

Our results emphasize that DETC and LC maturation occurs independently of microbiota and reciprocal immune cell interactions. However, we did not directly assess interactions with keratinocytes, which produce key cytokines and mediate cell–cell contacts critical for epidermal immune cell development. IL-34, for example, is primarily produced by keratinocytes during both development and steady state and supports LC differentiation (Wang *et al*, 2012; Greter *et al*, 2012), TGF-β1 can be derived from keratinocytes or produced via autocrine loops within LCs (Kaplan *et al*, 2007). Skint1, essential for the selection and maintenance of epidermal DETCs, is also predominantly expressed by keratinocytes (McKenzie *et al*, 2022; Barbee *et al*, 2011). While these findings underscore the role of niche-intrinsic cues in LC and DETC maturation, our study did not explicitly investigate keratinocyte–immune cell interactions. Thus, keratinocyte-derived signals may significantly contribute to the maturation of both cell types and warrant future investigation, ideally at single-cell resolution during peri- and postnatal development. Potential interactions between DETCs and LCs, allowing for coordinated postnatal maturation, have not yet been described. Based on our scRNA-seq and ligand–receptor inference analyses, we identified DETC-derived signals—namely *Il13*, *Csf2*, and *Tgfb1*—as candidate regulators of LC development and maturation. While TGF-β1 is well known to be essential for LC differentiation, DETCs have not previously been recognized as a source of this cytokine during epidermal development. IL-13 has been implicated in the regulation of dermal dendritic cell activation during steady state, but its potential role in LC maturation remains unexplored (Mayer *et al*, 2021). Similarly, CSF2 (GM-CSF) is required for *in vitro* LC differentiation, yet its function in postnatal LC maturation *in vivo* is poorly defined. (Geissmann *et al*, 1998; Herbst *et al*, 1998). Thus, future studies are needed to elucidate whether DETC-derived CSF2, IL-13, or TGF-β1 contribute functionally to LC maturation under physiological conditions.

In contrast, functional testing in *Tcrd^-/-^* mice—lacking γδ T cells—revealed no defects in LC maturation, as evidenced by an unchanged transcriptomic profile of LCs in the absence of γδDETCs in the adult epidermis. This observation may be explained by the compensatory presence of αβ T cells (αβDETCs) in *Tcrd^-/-^* mice, which likely preserve essential signaling cues required for LC differentiation. Supporting this, our cell-cell communication analysis showed that ligand–receptor interactions between αβDETCs and LCs closely mirrored those observed between γδDETCs and LCs, with no differences in interaction strength or ligand composition. Nevertheless, αβDETCs are known to decline from the epidermis after approximately three months of age (Jameson *et al*, 2004), raising the possibility that long-term LC homeostasis could be impaired in aging *Tcrd^-/-^* mice. Future studies in aged *Tcrd^-/-^* mice or *Cd3e^-/-^* mice, which lack all T cells, are needed to clarify the extent to which DETC–LC interactions are dispensable for both development and maintenance. Interestingly, our cell-cell interaction analysis identified LC-derived CCL4 as a potential signal toward DETCs. However, the predicted receptors on DETCs were CCR1 and CCR8, neither of which are known to directly bind CCL4 (Zlotnik & Yoshie, 2012). This suggests that LC-to-DETC communication may be limited or indirect. Further investigation, such as LC depletion studies during postnatal development, will be necessary to determine whether LCs have any functional role in DETC maturation.

Our results further highlight that the maturation of the entire epidermal immune compartment is not affected by either the absence or increased complexity of microbiota during postnatal development. In other organs, the maturation of tissue-resident macrophages has been shown to depend on microbial cues and microbiota-derived metabolites (Kolter *et al*, 2019; Thion *et al*, 2018). By contrast, other barrier tissues, such as the oral mucosa, exhibit dramatic microbiota-dependent differences in immune cell composition, ontogeny, and turnover (Capucha *et al*, 2018). Whether commensal metabolites are excluded from the epidermal niche by the perinatally established basal membrane, or whether such metabolites are accessible but redundant for epidermal immune development, remains unresolved. Notably, DETCs and LCs have been implicated in tolerance induction to skin commensals (Gribonika *et al*, 2025; Love-Schimenti & Kripke, 1994), and LCs have been reported to scavenge microbiota and antigens from the skin surface by extending dendrites through the tightly connected keratinocyte layers (Nishibu *et al*, 2006). Despite these established functions, our data demonstrate that their own maturation is likely governed by tissue-intrinsic programs, with microbiota-derived signals appearing dispensable for this process. Since our study focused on the steady-state differentiation dynamics and interactions of DETCs and LCs, we cannot exclude the possibility that their interactomes during perinatal development contribute to the establishment of tissue immunity and commensal tolerance. Future studies are needed to investigate how the absence of DETCs or LCs affects immune responses to pathogens, tolerance to skin commensals, and the induction of self-tolerance during early life.

In summary, our study delineates developmental programming of DETCs and LCs in the early-life mouse epidermis. By uncovering their potentially independent differentiation trajectories and highlighting the dispensability of microbiota for their development, we establish a foundational resource for future investigations into skin immune homeostasis and early-life tissue education.

## MATERIALS AND METHODS

### Mice

*C57BL/6J* mice from Charles River were used as wildtype mice. All mice were housed in specific pathogen-free conditions and backcrossed on *C57BL/6J* genetic background, if not otherwise stated in the manuscript. Animals were kept in a 12h/12h dark-light cycle and water and food was provided *ad libitum*. All animal experiments were approved by local administration (Regierungspräsidium Freiburg) and were performed in accordance with the respective national and institutional regulations. Transgenic mouse lines including *Tcrd^-/-^*(*B6.129P2-Tcrdtm1Mom/J*) (Itohara *et al*, 1993) and *Cx3cr1^GFP^* (*B6.129P2(Cg)-Cx3cr1tm1Litt/J* (Jung *et al*, 2000) were used in this study. Wildlings (*C57BL/6NTac)* were created by embryo transfer of laboratory mice into pseudopregnant wild mice (Rosshart *et al*, 2019). Germfree animals were obtained from the Clean Mouse Facility (Bern, Switzerland). These mice were born and raised in a completely sterile milieu in pressurized HEPA-filtered plastic film isolators (Smith *et al*, 2007).

### Timed mating

Timed mating was used to analyze embryos at defined developmental stages. The matings were started in the evening and separated after vaginal plug check on the following morning. Embryonic development was estimated considering the day of vaginal plug formation as embryonic day (E) 0.5 and staged by developmental criteria (Theiler, 1972). Embryos were dissected at E17.5 and E18.5. Postnatal time points P0, P1, P2, P4, P7, P14, P21, P30, P60 and P90 and P100 were dissected referring to the day of birth.

### Dissection of peri- and postnatal epidermis

Pregnant mice were euthanized with CO_2_ followed by cervical dislocation. Embryos at E17.5 and E18.5 as well as neonates from P0-P7 were decapitated and ventral skin was dissected. For P4 and P7, hair was removed from abdominal skin using depilatory cream and mechanical detachment after 5 minutes (min). Skin samples were washed in ice-cold 1x PBS. The skin was digested for 45 min at 37°C in enzyme mix (Dispase II (2,4 mg/ml), 3% FCS in 1x PBS). Afterwards, the epidermis was carefully separated from the dermis using fine forceps.

### Dissection of adult epidermis

Mice at the age of P14 onwards were euthanized with CO_2_ followed by cervical dislocation. Skin tissue from mice at P14 and older was obtained from ears. Hair was removed from ears with depilatory cream and mechanical detachment after 5 min. Skin samples were washed in ice-cold 1x PBS. Ventral and dorsal sheets of the ears were separated. The skin was digested for 45 min at 37°C in enzyme mix 1 (Dispase II (2,4 mg/ml), 3% FCS in PBS). Afterwards, epidermis was carefully separated from the dermis using fine forceps.

### Immunofluorescence imaging of whole mount epidermis

Neonatal ventral epidermis was fixed for 2 hours, and ear epidermis was fixed for 20 min in 4% PFA at 4°C. Epidermal samples were washed three times with 1x PBS and blocked for 2 hours at 4°C with 1 ml of blocking buffer 1 (0.5 % BSA, 5 % normal goat serum, 0.3 % Triton X-100 in 1x PBS). Epidermis was then incubated overnight at 4°C with primary antibody mix (anti-Langerin 1:100 [clone eBioRMUL.2, rat anti-mouse, eBioscience]; anti-EPCAM 1:300 [clone G8.8, rat anti-mouse, ThermoFisher]; anti-F4/80 1:300 [clone EPR2645-166, chicken anti-mouse, abcam]; anti-CD3e 1:300 [clone SP162, rabbit anti-mouse, abcam]; and anti-Ki67 1:500 [clone SP6, chicken anti-mouse, abcam and clone SolA15, rat anti-mouse, ThermoFisher]) in blocking buffer 2 (0.5 % BSA, 0.3 % Triton X-100 in 1x PBS). After rinsing three times with blocking buffer 2, the secondary antibody mix (donkey anti-rat Alexa Fluor® 488 1:300; goat anti-chicken Alexa Fluor® 488 1:300; goat anti-rabbit Alexa Fluor® 568 1:300; donkey anti-rabbit Alexa Fluor® 647 1:300; goat anti-rat Alexa Fluor® 647 1:300; DAPI 1:5000) was added and incubated for 2 h at room temperature. Sheets were washed again three times with blocking buffer 2 and subsequently mounted using Mowiol mounting medium. LC and DETC numbers during development were quantified by confocal imaging with a SP8 X with white-light laser (Leica) using a 20x objective. Representative close-up images of LC and DETC development were taken using a 63x objective. Ki67 quantification was performed using the Zeiss LSM710 confocal microscope using a 20x objective and Z-stacks of 0.2 to 1 µm. We analyzed Z layers separately to identify Ki67^+^ DETCs and LCs and distinguish them from proliferating basal stem cells in the epidermis. For imaging of DETCs and LCs in wildling samples, a Leica Thunder Imager with a 20x objective was used. Representative images of epidermal sheets from wildlings and SPF animals were taken with the Zeiss LSM710 Confocal using a 20x objective. Post-acquisition editing and quantification of all representative figures was done with the Fiji platform (Schindelin *et al*, 2012).

### scRNA-seq of the epidermal tissue

Isolated epidermis was incubated for 20 min at 37°C in enzyme mix 2 (Collagenase-D (1 mg/ml), DNase (0,2 mg/ml), 3% FCS in PBS). Epidermis was mechanically homogenized through a 100 μm cell strainer into FACS buffer (0,5 % BSA, 2 mM EDTA in PBS). Tubes for the collected cells were pre-coated with 10 % BSA for 2 h. Cells were centrifuged at 320 g for 7 min at 4°C and resuspended in Fc receptor blocking antibody anti-CD16/32 (clone 2.4G2, BD Biosciences, 1:200) for 20 min on ice to prevent non-specific antibody binding. Afterwards, cells were incubated with anti-Gr1 (clone RB6-8C5, BD Biosciences, 1:200) and anti-CD19 (clone 6D5, BioLegend, 1:200) to exclude granulocytes and B cells and an anti-CD45 (clone 30-F11, eBioscience, 1:200) antibody to gate on DETCs and LCs. A fixable viability dye (eBioscience, 1:1000) was used to exclude dead cells. Additionally, 1 μl of hashing antibodies was added per sample. Hashing antibodies were obtained as purified and already oligo-conjugated in TotalSeq-C (5’ chemistry) format from BioLegend. Cells were stained for 20 min and washed. CD45^+^ immune cells were sorted using a BD FACSAria^TM^ III, a BD FACSAria^TM^ Fusion or a BC MoFlo Astios EQ in the Lighthouse Core Facility, University of Freiburg. Sorted cells were processed through the 10x Genomics single-cell 5’ v2 workflow according to manufacturer’s instructions. Libraries were pooled to desired quantities to obtain appropriate sequencing depths as recommended by 10x Genomics and sequenced on a NovaSeq6000 sequencer.

### Quantification of transcript abundance and downstream scRNA-seq data analysis

Gene expression and hashtag abundance were quantified using the count command from cellranger-6.0.0 and cellranger-7.2.0 using the prebuilt CellRanger mouse mm10 references from 2020. The scRNA-seq data was analyzed using python (v3.12.4) and the packages scanpy (v1.10.4) (Wolf *et al*, 2018) and muon (v0.1.6) (Bredikhin *et al*, 2022). The hashtags were demultiplexed using hashsolo (Bernstein *et al*, 2020) as included in scanpy and batch effects were removed using Harmony (Korsunsky *et al*, 2019). Cells containing less than 500 counts or 200 genes, or more than 10% mitochondrial counts were removed as low-quality cells. Genes that appeared in less than three cells were also removed. Importantly, ribosomal, and mitochondrial genes, as well as predicted genes with the *Gm*-identifier, were excluded from the analysis after the quality control. The counts were normalized and set to a target sum of 10000, followed by log1p transformation and scaling with a cutoff at 10. To determine the top 2000 highly variable genes the flavor “seurat_v3” was used in the highly_variable_genes function. Dimensionality reduction was performed using uniform manifold approximation and projection (UMAP) while making sure that the neighborhood graph was calculated on the Harmony corrected PCA embedding. Clustering was performed using the leiden algorithm and clusters were annotated based on marker genes found after calculating differentially expressed genes. For downstream analysis of DETCs only canonical Vγ5Vδ1 DETCs (expressing *Tcrg-V5* and *Trdv4*) were kept.

### Trajectory analysis of scRNA-seq data

In order to include temporal information in the trajectorial analysis, a temporal problem from Moscot (Schiebinger *et al*, 2019; Klein *et al*, 2025) was built and passed to CellRank (Lange *et al*, 2022; Weiler *et al*, 2024), to build a real-time kernel. Additionally, a connectivity kernel was built using the neighborhood graph. These were combined into a single kernel. Then a generalized probabilistic canonical correlation analysis (GPCCA) estimator was created using the combined kernel and initial and terminal states were predicted. Since multiple terminal states were predicted, the trajectory leading to the terminal state closest to our last time point was picked as the differentiation trajectory of interest. The initial state was used to determine the root cell for diffusion pseudotime calculation based on a PAGA graph at cluster resolution. DPT was then used as the trajectory to smooth the expression of genes. Smoothed gene profiles were used for clustering by building an AnnData object and clustering using the Leiden algorithm as implemented in CellRank and scanpy.

## Cell-cell communication inference from scRNA-seq data

Cell-cell communication was inferred using the packages LIANA and Tensor-cell2cell (Dimitrov *et al*, 2024; Armingol *et al*, 2022). For this purpose, the resource “mouseconsensus” from LIANA was used in combination with the methods logfc, geometric_mean, singlecellsignalr, connectome, cellphonedb, natmi, and cellchat. Subsequent analysis was performed using a consensus score from all these methods as implemented in LIANA. These methods were used to calculate a consensus score for all replicates separately and the results were used to build a 4D tensor using Tensor-cell2cell. This tensor was then decomposed into several factors, which was picked according to package instructions.

## ACKNOWLEDGMENTS

We thank Maria Oberle for excellent technical assistance. We thank the Lighthouse Core Facility (LCF) for their support with cell sorting. LCF is funded in part by the Medical Faculty, University of Freiburg (Project Numbers 2021/A2-Fol; 2021/B3-Fol) and the Deutsche Forschungsgemeinschaft (DFG, German Research Foundation) through Project Number 450392965. S. is supported by the Department of Medicine II, Freiburg University Medical Center, Faculty of Medicine, University of Freiburg. This study was supported by the DFG through TRR 359 (Project ID 491676693 to J.K., D.E., M.P., P.H., S.P.R.., K.K. and S.), SFB 1160 (Project ID 256073931 to M.P., P.H., S.M., S.P.R.., K.K. and S.), 322977937/GRK2344 (S.), TRR167 (Project ID 259373024 to D.E., M.P. and K.K.), CRC1479 (Project ID 441891347 to M.P., S.M., and K.K.) and by the DFG under Germany’s Excellence Strategy (CIBSS—EXC-2189, Project ID 390939984 to M.P., S.M., W.W.S., P.H. and K.K.). K.K. is supported by the Heisenberg program of the DFG (Project ID 544402801). D.E. was further supported by the Else Kröner-Fresenius Foundation and the Ministry of Science, Research and Arts, Baden-Wuerttemberg under the aegis of JPND. S.P.R. was supported by the DFG Emmy Noether-Programm RO 6247/1-1 (Project ID 446316360), and the TRR 417 (Project ID 540805631).

## AUTHOR CONTRIBUTIONS

Conceptualization, S., K.K.; Methodology, S. and K.K.; Formal Analysis, D.O., A.O., L.M.K., S., K.K.; Investigation, D.O., A.O., L.M.K., C.C., S.D., K.B., S.R., N.G., M.E.; Resources, V.F., D.E., J.K., S.M., W.W.S., M.P., P.H., S.P.R..; Writing, D.O, A.O., L.M.K., S., K.K.; Visualization, D.O., A.O., L.M.K.; Supervision, S., K.K.; Project Administration, S., K.K.; Funding Acquisition, S., K.K.

## DATA AND MATERIALS AVAILABILITY

The primary read files and the raw counts as well as processed data for all single-cell RNA sequencing datasets reported in this paper will be available to download from the Gene Expression Omnibus platform upon publication. Processed data can be downloaded from https://github.com/D-Obwegs/DETC-and-LC-development-scRNA-seq. Codes to reproduce the data analysis and figures are available at: https://github.com/D-Obwegs/DETC-and-LC-development-scRNA-seq.

---

**Supp. Figure 1.**
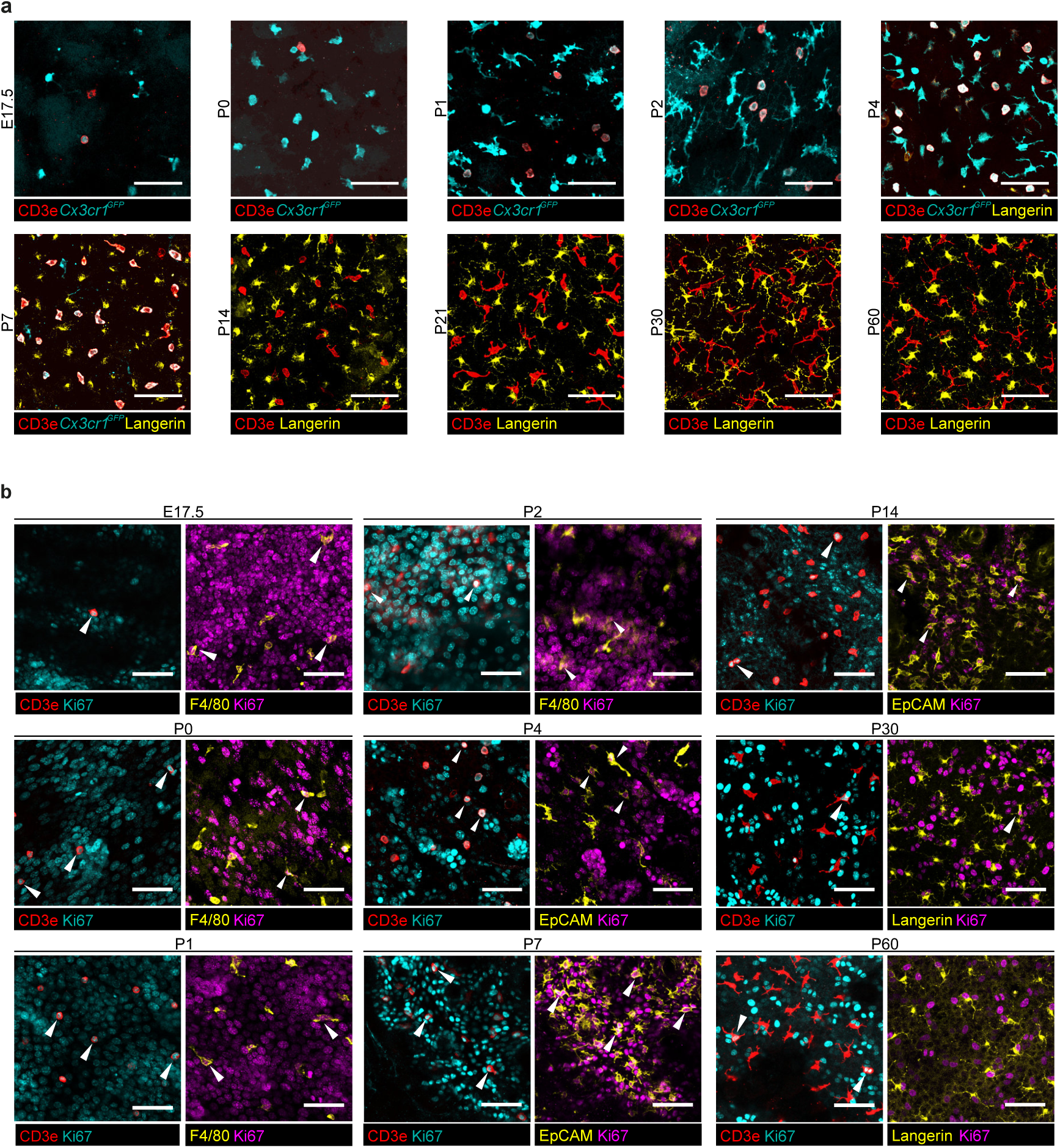
Epidermal networks of DETCs and LCs expand from birth to adulthood. **a** Representative whole mount confocal images of epidermal DETCs and LCs at all analyzed time points. CD3e is shown in red, *Cx3cr1^GFP^* is shown in cyan, and Langerin is shown in yellow. Maximum projection of the confocal z-stack is shown. Scale bar= 50 µm. **b** Representative whole mount confocal images of epidermal Ki67^+^ DETCs and LCs at all analyzed time points. CD3e is shown in red, Ki67 is shown in cyan or magenta, and F4/80, EpCAM or Langerin are shown in yellow. Arrowheads indicate proliferating DETCs or LCs. Selected z layers are shown for illustration purposes. Scale bar= 50 µm.

**Supp. Figure 2.**
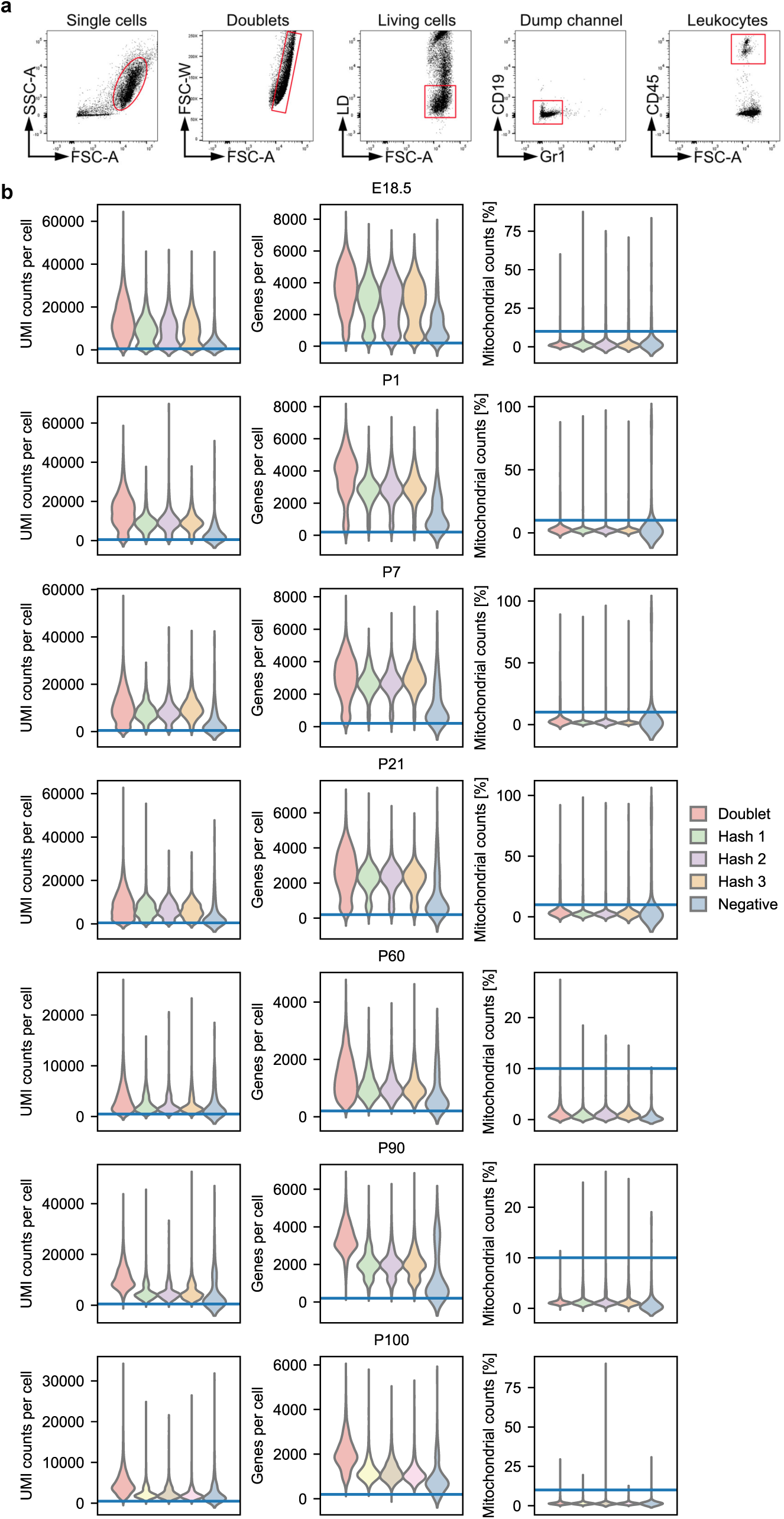
Epidermal immune cell isolation and quality control of scRNA-seq data. **a** Representative flow cytometry plots showing the gating strategy used for single cell sorting of immune cells from epidermal tissues. **b** Violin plots depicting the number of UMI counts (left), number of genes (middle), and the percentage of mitochondrial genes (right) per cell after demultiplexing separated by assignment to either a hashing antibody, doublet or negative. Color is indicated in legend.

**Supp. Figure 3.**
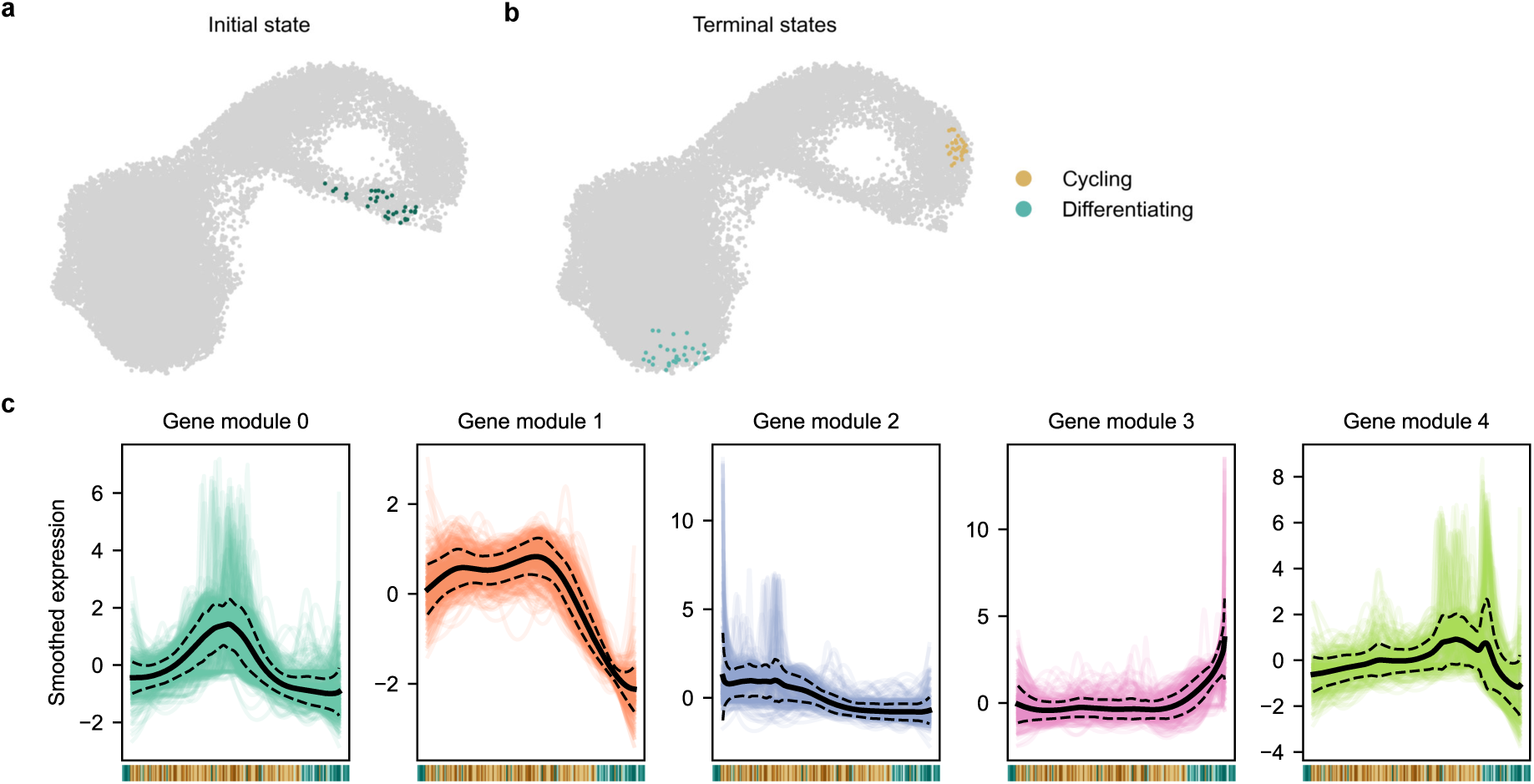
Trajectory analysis of DETCs across development. **a** UMAP plot showing the initial state (green) as predicted by the package CellRank. **b** UMAP plot showing the terminal state (cycling (orange) and differentiated (cyan)) as predicted by the package CellRank. **c** Gene modules as identified by Leiden clustering of smoothed gene expression of top 2000 highly variable genes along pseudotime. Continuous line represents the mean of all genes in the module and dashed lines represent the mean ± standard deviation. Colors represent the respective modules as shown in Fig. 3 h, i.

**Supp. Figure 4.**
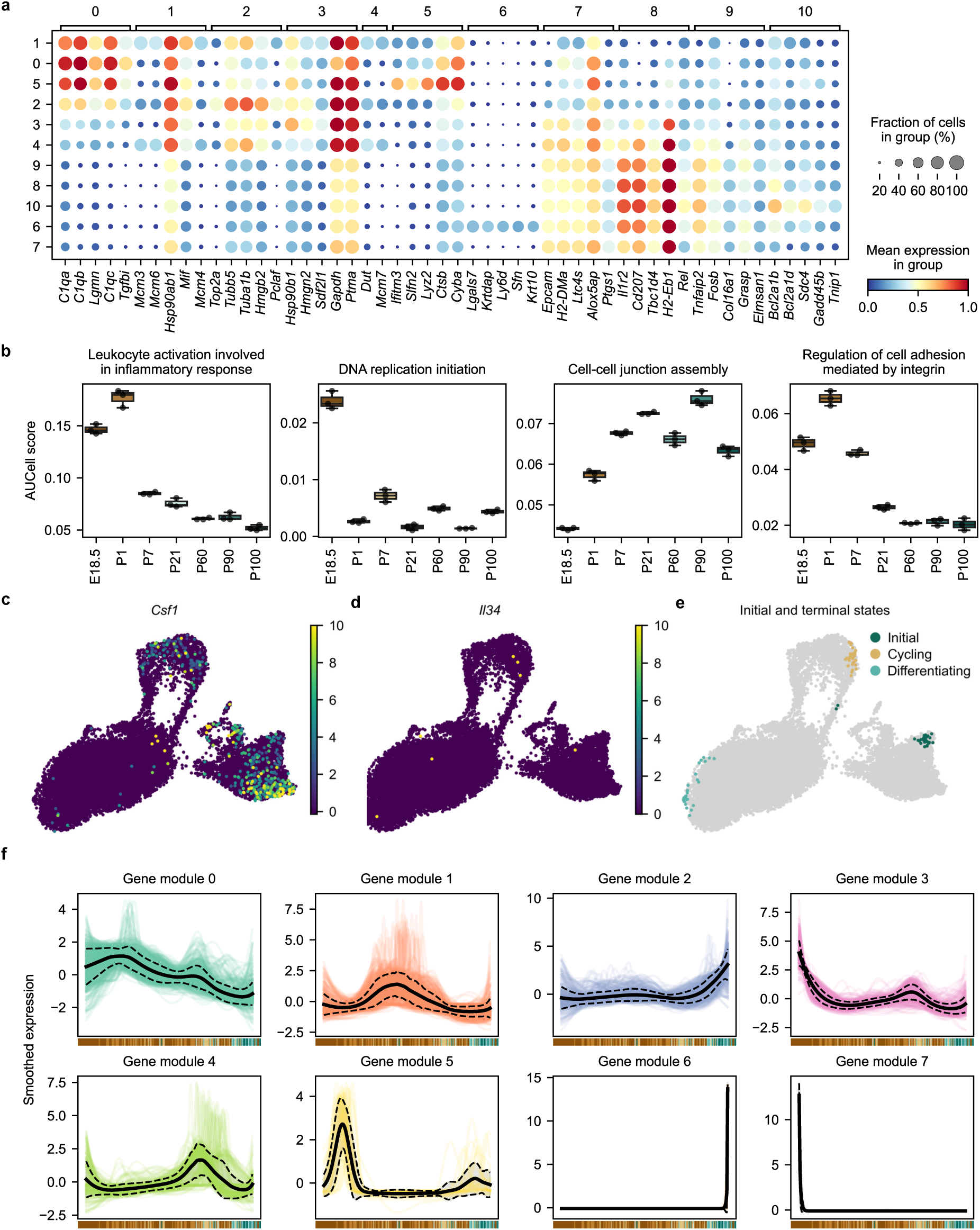
LCs employ distinct gene programs during their postnatal maturation. **a** Dot plot of top 5 DEGs per cluster. Color represents scaled mean expression of the gene in the respective cluster and dot size represents the fraction of cells expressing the gene. Cluster number is indicated in legend. **b** Box plot showing mean AUCell scores per replicate of selected GO terms across the different time points. Mean +/- SEM is shown. Dots represent individual biological samples. **c** Expression of *Csf1* on UMAP embedding based on gene expression profiling. Scale reflects the relative gene expression. **d** Expression of *Il34* on UMAP embedding based on gene expression profiling. Scale reflects the relative gene expression. **e** UMAP plot depicting the initial (green), terminal (cycling (orange) and differentiating (cyan) states as predicted by CellRank indicated by color. **f** Gene modules as identified by Leiden clustering of smoothed gene expression of top 2000 highly variable genes along DPT. The continuous line represents the mean of all genes in the module and dashed lines show the mean ± standard deviation. Colors represent the respective modules as shown in Fig 4 h, i.

**Supp. Figure 5.**
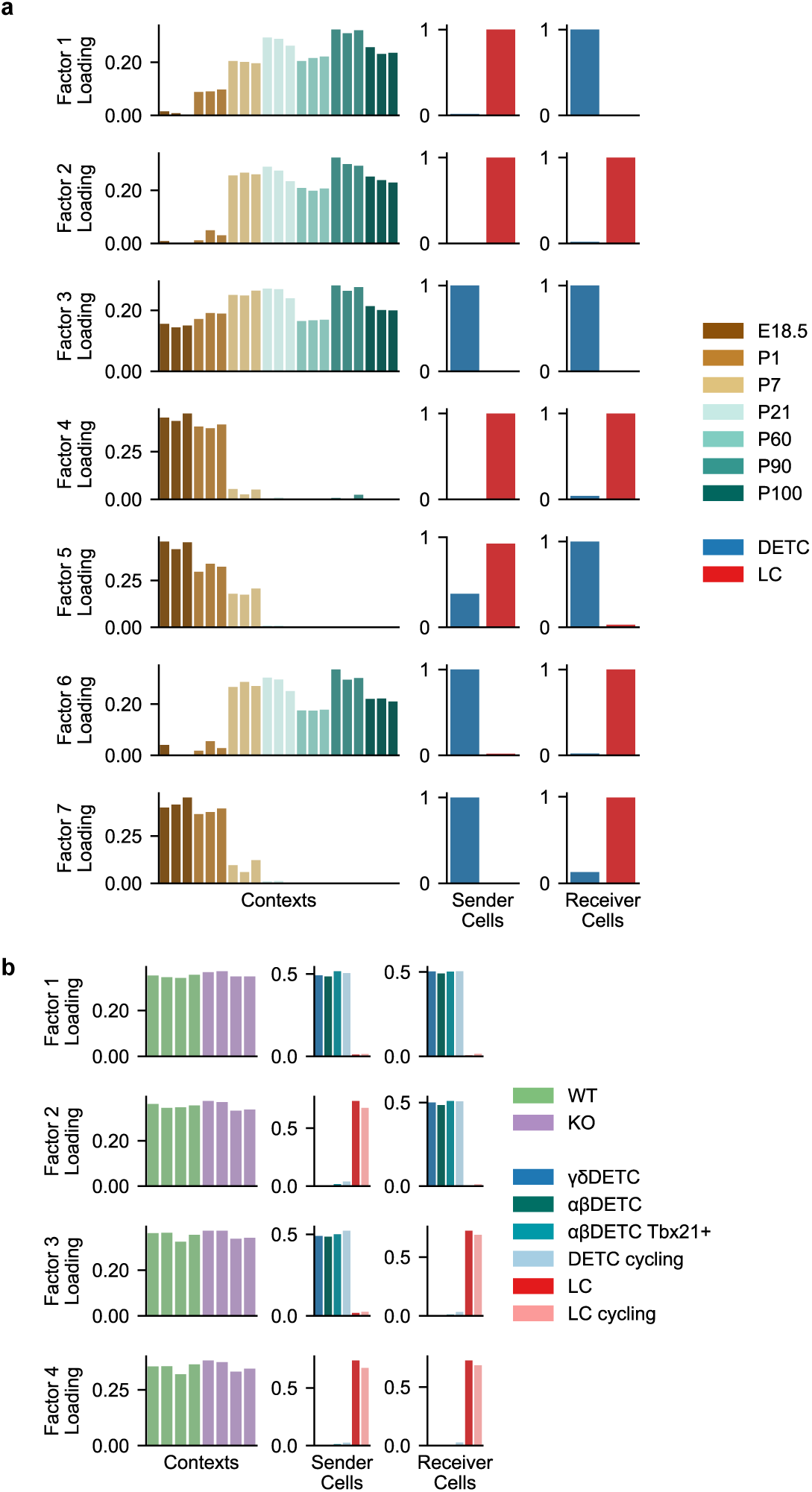
DETCs and LCs display a defined and concise interactome across development. **a** Left: Graphs depicting results of running Tensor-cell2cell frameworks on our dataset containing different time points. Each row represents a factor displaying paracrine or autocrine signaling between DETCs (blue) and LCs (red), and each column a tensor dimension, wherein each bar plot represents an element of that dimension (time point, a sender cell or a receiver cell). Factor loadings (y-axis) are displayed for each element of a given dimension. Colors of individual time points are depicted in legend. Right: Scheme (showing autocrine and paracrine signaling between DETCs and LCs of the factors 1-7 in early (top-right) and adult (bottom-right) time points. **b** Graphs depicting results from running Tensor-cell2cell frameworks using the αβ and γδDETCs and LCs from *Tcrd^+/+^* (WT) and *Tcrd^-/-^* (KO) mice. Each row represents a factor displaying paracrine or autocrine signaling, and each column a tensor dimension, wherein each bar plot represents an element of that dimension (genotype, sender, or receiver cells). Factor loadings (y-axis) are displayed for each element of a given dimension. Colors are indicated in legend.

**Supp. Figure 6.**
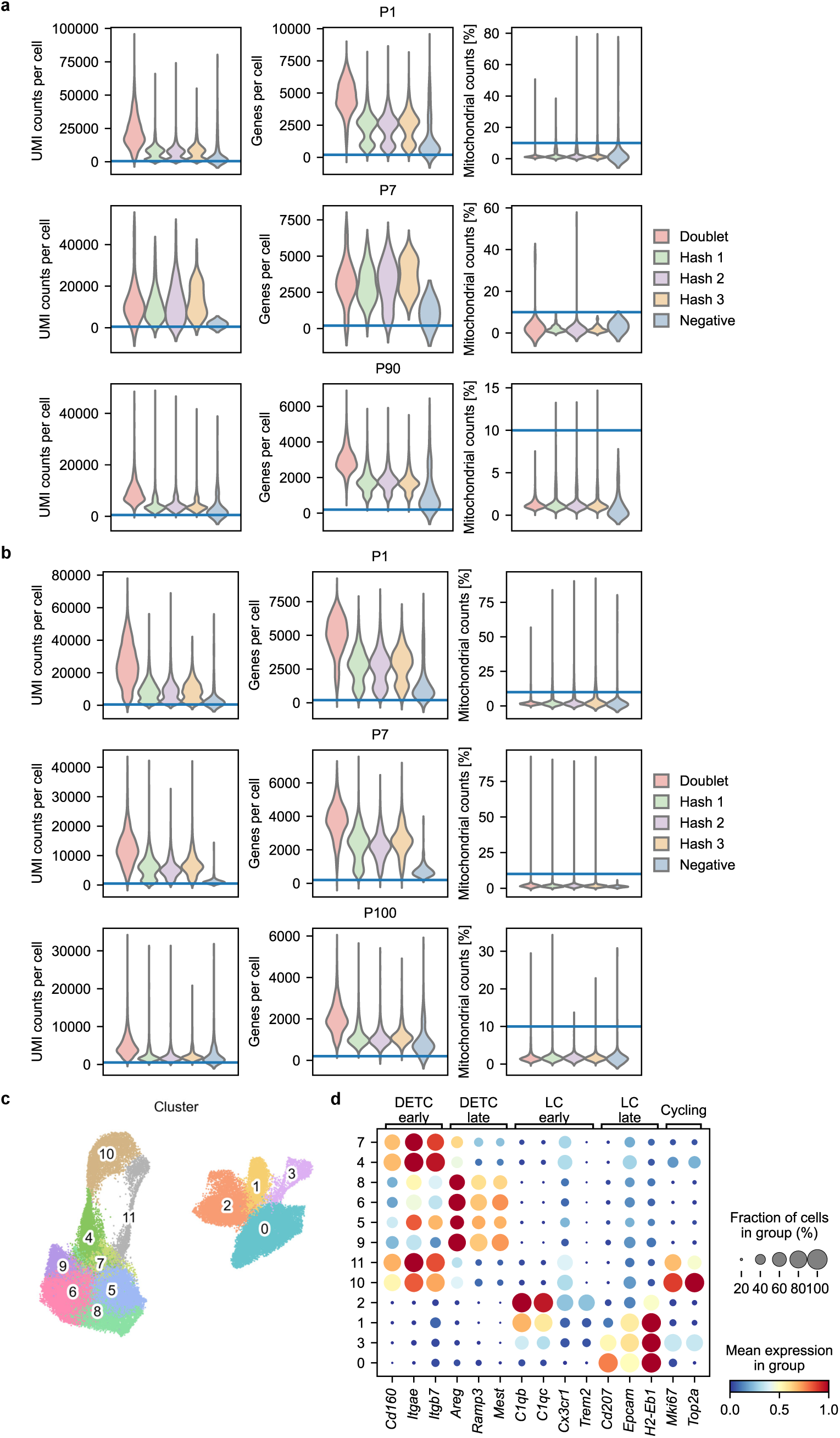
Microbial colonization of the skin does not affect DETC and LC maturation. **a** Violin plots depicting the number of UMI counts (left), number of genes (middle), and the percentage of mitochondrial genes (right) per cell after demultiplexing for germfree mice separated by assignment to either a hashing antibody, doublet or negative. Color is indicated in legend. **b** Violin plots showing the number of UMI counts (left), number of genes (middle), and the percentage of mitochondrial genes (right) per cell after demultiplexing for wildlings separated by assignment to either a hashing antibody, doublet or negative. Color is indicated in legend. **c** UMAP plot showing 12 identified cell clusters of GF mice, SPF mice and wildlings (N = 80556 cells). Each color represents one cell cluster. **d** Dot plot showing expression of selected genes in the 12 different clusters for early and late DETCs/LCs and cycling cells. Color represents the scaled mean expression of the gene in the respective cluster and dot size represents the fraction of cells in the cluster expressing the gene.

**Supp. Figure 7.**
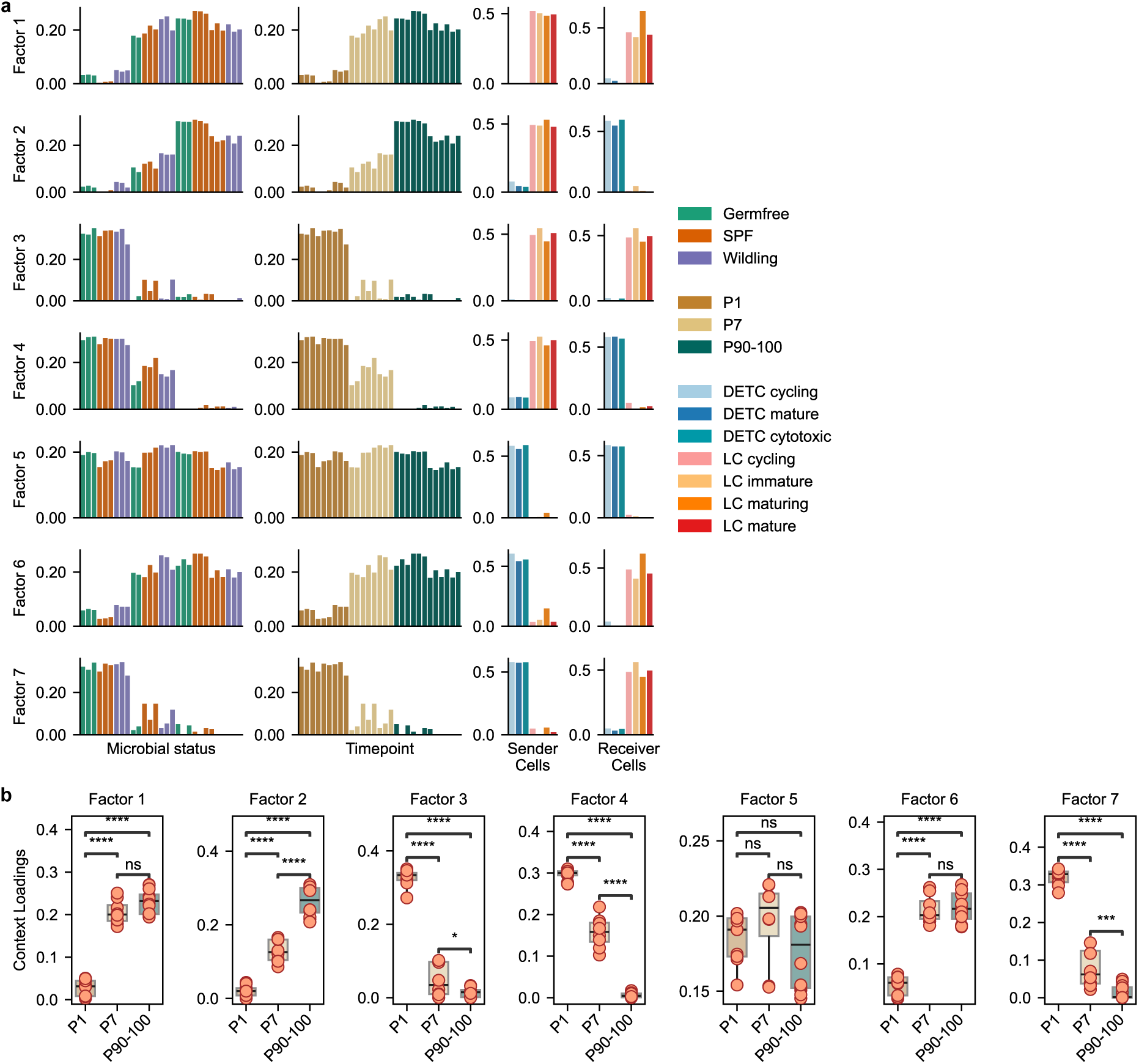
Developmental time point, rather than microbial status defines DETC and LC maturation. **a** Graphs depicting results of running Tensor-cell2cell frameworks on our dataset containing different microbial statuses, time points and cell states (cycling, immature, maturing, mature and cytotoxic). Each row represents a factor, and each column a tensor dimension, wherein each bar plot represents an element of that dimension (microbial status, time point, a sender cell or a receiver cell). Factor loadings (y-axis) are displayed for each element of a given dimension. Color is indicated in legend. **b** Box plots of the context loadings for all seven factors recovered by factor decomposition of a tensor built using microbial status and time points as contexts. Box plots show the median (line), interquartile range (box), and 1.5x interquartile range (whiskers). Statistical testing was performed using Mann-Whitney U rank test with Benjamini-Hochberg correction for multiple testing.

